# A multi-dimensional integrative scoring framework for predicting functional variants in the human genome

**DOI:** 10.1101/2021.01.06.425527

**Authors:** Xihao Li, Godwin Yung, Hufeng Zhou, Ryan Sun, Zilin Li, Yaowu Liu, Iuliana Ionita-Laza, Xihong Lin

**Affiliations:** Department of Biostatistics, Harvard T.H. Chan School of Public Health, Boston, MA 02115, USA; Genentech/Roche, South San Francisco, CA 94080, USA; Department of Biostatistics, University of Texas MD Anderson Cancer Center, Houston, TX 77030, USA; School of Statistics, Southwestern University of Finance and Economics, Chengdu, Sichuan, China; Department of Biostatistics, Columbia University Mailman School of Public Health, New York, NY 10032, USA; Program in Medical and Population Genetics, Broad Institute of Harvard and MIT, Cambridge, MA 02142, USA; Department of Statistics, Harvard University, Cambridge, MA, 02138, USA

## Abstract

Attempts to identify and prioritize functional DNA elements in coding and noncoding regions, particularly through use of in silico functional annotation data,continue to increase in popularity. However, specific functional roles may vary widely from one variant to another, making it challenging to summarize different aspects of variant function. Here we propose Multi-dimensional Annotation Class Integrative Estimation (MACIE), an unsupervised multivariate mixed model framework capable of integrating annotations of diverse origin to assess multi-dimensional functional roles for both coding and noncoding variants. Unlike existing one-dimensional scoring methods, MACIE views variant functionality as a composite attribute encompassing multiple characteristics, and estimates the joint posterior functional probability vector of each genomic position, a quantity that offers richer and more interpretable information in the presence of multiple aspects of functionality. Applied to a variety of independent coding and non-coding datasets, MACIE demonstrates powerful and robust performance in discriminating between functional and non-functional variants. We also show an application of MACIE to fine-mapping using lipids GWAS summary statistics data from the European Network for Genetic and Genomic Epidemiology Consortium.

## Introduction

Ever since the completion of the human genome sequence, substantial effort has been invested into identifying and annotating functional DNA elements. For any given genetic variant, a diverse set of functional annotations is now available. For example, the computational tool PolyPhen^1^ predicts damaging effects of missense mutations. PhastCons^2^, PhyloP^3^, and GERP++^4^ leverage comparative sequence information to identify regions that show evolutionary conservation. The Encyclopedia of DNA Elements (ENCODE) has extensively mapped regions of transcription factor binding, chromatin structure, and histone modification, effectively assigning biochemical functions for ∼80% of the genome^5^. Other initiatives such as the Roadmap Epigenomics project^6^ and FANTOM5 project^7,8^ also provide evidence for potential regulatory variants in the human genome.

Although functional annotations vary considerably with respect to the specific elements they evaluate and the extent of the human genome they annotate, it is well understood that they provide complementary lines of evidence^9^. Therefore, in order to obtain a comprehensive understanding of the biological relevance of genomic segments, all of the information provided by different annotations should be jointly synthesized. However, it remains challenging how to summarize these diverse functional annotations in an insightful and interpretable manner.

Current algorithmic scoring frameworks utilize a variety of statistical and machine-learning methods to aggregate information from large, diverse sets of individual annotations into single measures of functional importance. Supervised tools such as CADD^10^, DANN^11^, GWAVA^12^, FATHMM-MKL^13^, and FATHMM-XF^14^ build machine learning classifiers on training sets with pre-labeled functional statuses, e.g., fine-mapped pathogenic or disease-associated variants labeled against benign or neutral variants. Such supervised approaches rely strongly on the quality of labels in the training set. Therefore, they may demonstrate suboptimal performance when inaccurate or biased labels are used. Unsupervised methods such as EIGEN^15^, GenoCanyon^16^, PINES^17^, and FUN-LDA^18^ do not rely on any labeled training data. They possess advantages in studying non-coding regions, where our current lack of knowledge often precludes gold-standard training data labels. A third group of methods including fitCons^19^ and LINSIGHT^20^ use evolution-based approaches that characterize the potential effect of natural selection at each genomic location using polymorphism and divergence data. Recent reviews provide a more detailed discussion of available functional annotation tools^21,22^.

Although existing methods attempt to integrate functional annotations through various approaches, to the best of our knowledge, these methods all summarize the annotation information with a single rating. In doing so, they implicitly assume that variant function can be described along a single axis, with variants being more functional on one end of the axis and less functional on the other end. This assumption may be reasonable if interest lies in predicting a specific aspect of variant function (e.g. regulatory behavior) and all annotations used as input are intended to predict that same aspect. However, if multiple aspects of variant function are simultaneously of interest, then it is unclear how to interpret the one-dimensional consolidation of annotations measuring different aspects of function, especially when these annotations appear to provide orthogonal information, e.g., weak correlation between evolutionary conservation scores and regulatory scores (**Figure 1**). Therefore, it is of interest to construct multi-dimensional integrative scores capable of capturing multiple facets of variant function simultaneously.

**Figure 1.**
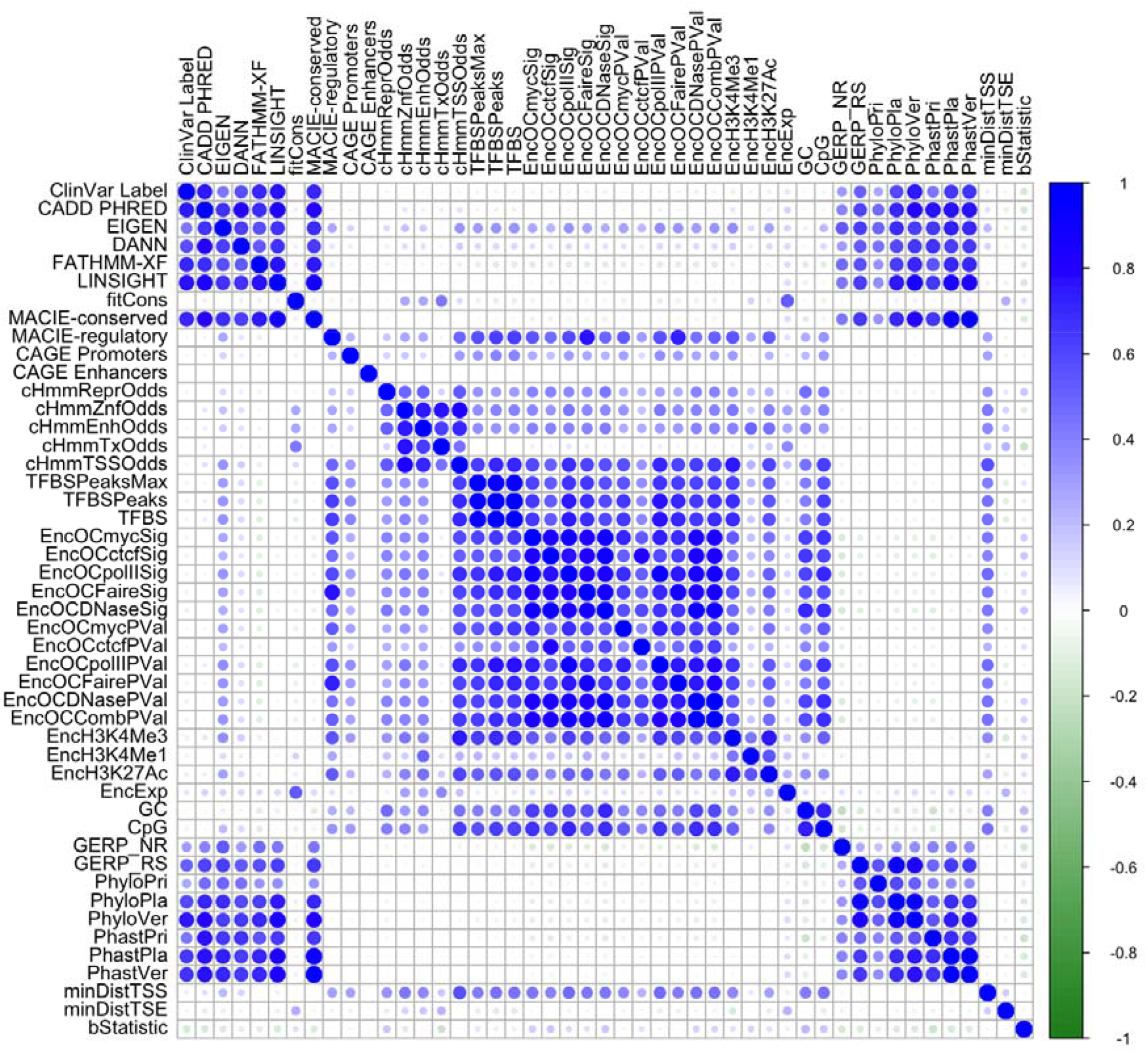
Heatmap demonstrating the correlation between individual and integrative functional scores for ClinVar pathogenic and benign noncoding variants.

In this paper we propose Multi-dimensional Annotation Class Integrative Estimation (MACIE), an unsupervised multivariate mixed model framework capable of synthesizing multiple categories of annotations and producing interpretable multi-dimensional integrative scores (**Supplementary Figure 1**). Instead of a single rating, MACIE explicitly defines variant function as a vector of binary outcomes, each outcome capturing functionality corresponding to a specific class of annotations. Correlations within and between the different classes of annotations are explicitly modeled, another advancement over existing methods. Using the Expectation-Maximization algorithm, MACIE calculates the joint posterior probability vector of a genomic position being functional (**Methods**).

Because of its multivariate formulation, MACIE is able to provide detailed and nuanced assessments of variant functionality. Output from MACIE is highly interpretable due to the specificity allowed by multiple functional classes.

Additionally, the MACIE framework allows for considerable versatility to incorporate data in a manner that is most biologically relevant to the scientific question of interest. We apply MACIE to multiple independent coding and noncoding testing sets and show that, compared to current state-of-the-art integrative scores, MACIE consistently provides robust and best or near best performance in discriminating between functional and non-functional variants.

## Results

### Construction of MACIE training sets

MACIE scores were computed for (1) nonsynonymous coding and (2) noncoding and synonymous coding variants separately because the two types of variants are expected to have highly different functional profiles^15^. All nonsynonymous coding annotations and some noncoding and synonymous coding annotations were downloaded from EIGEN. The remaining noncoding and synonymous coding annotations were downloaded from CADD full database^10^ v1.3.

### Nonsynonymous coding variants

For the nonsynonymous coding training set, we randomly extracted 10% of the variants with a match in the dbNSFP database^23^. This database excludes synonymous variants that fall in coding regions but do not alter protein function. Only one unique variant per position was selected, and variants residing in sex chromosomes X and Y were removed to mitigate potential sources of bias. The final set included approximately 2.2 million variants. For each variant in the training set, four protein substitution damage scores (SIFT^24^, PolyPhenDiv, PolyPhenVar^1^, Mutation Assessor^25^) and eight evolutionary conservation scores (GERP_NR and GERP_RS^4^; PhyloP primate (PhyloPri), placental mammal (PhyloPla), and vertebrate (PhyloVer)^3^; PhastCons primate (PhastPri), placental mammal (PhastPla), and vertebrate (PhastVer)^2^) were extracted from the EIGEN database^15^. Thus we defined the two-class MACIE model *(M=2)* for nonsynonymous coding variants to assess damaging protein coding function and evolutionarily conserved function. Full information on the MACIE model for nonsynonymous coding variants and the list of individual functional scores are given in **Methods** and **Supplementary Table 1**.

### Noncoding and synonymous coding variants

For the noncoding and synonymous coding training set, we extracted a random sample comprising 10% of the variants in the 1000 Genomes Project dataset that were located within 500 bp upstream of a gene start site and did not possess a match in dbNSFP. Duplicated variants with multiple alternative alleles and variants in sex chromosomes X and Y were again removed to mitigate potential bias. The final training set included 36,431 variants. For each variant in the training set, the same eight evolutionary conservation scores used for coding variants were extracted from the EIGEN full database^15^. A total of twenty-eight transformed epigenetic scores were additionally extracted from the CADD database^10^ v1.3, including a collection of regulatory annotations from the ENCODE Project^5^, three transcription factor binding site scores, GC content, CpG content, five chromatin state probabilities derived from the 15 state ChromHMM model^26^, a background selection score^27^, and physical distance metrics^10^. We then defined the two-class MACIE model *(M=2)* for noncoding and synonymous coding variants to assess evolutionarily conserved function and epigenetic regulatory function. Full information on the MACIE model for noncoding and synonymous coding variants and the list of individual functional scores are given in **Methods** and **Supplementary Table 1**. Detailed information on pre-processing steps for the epigenetic scores are given in **Supplementary Table 2**.

### Benchmarking the performance of MACIE with other integrative scoring methods

We compared the predictive performance of MACIE against existing state-of-the-art variant classifiers including CADD^10^, FATHMM-XF^14^, EIGEN^15^, fitCons^19^, LINSIGHT^20^, and DANN^11^ over a range of realistic variant assessment scenarios. Specifically, we assessed the ability of each score to identify clinically significant variants from ClinVar^28,29^; loss-of-function variants in the *BRCA1* gene uncovered through saturation genome editing (SGE)^30^; promoters and enhancers from the FANTOM5 project defined by cap analysis of gene expression (CAGE)^7,8^; and experimentally verified functional variants from massive parallel reporter assays (MPRA)^31,32^. Some alternative scoring methods were excluded due to difficulties related to providing a proper comparison of results. For example, LINSIGHT is designed to predict the deleteriousness of noncoding variants, so we did not include it in the comparison for nonsynonymous coding variants.

### Distribution of posterior probabilities for noncoding and synonymous coding variants in the training set

In **Supplementary Table 3** we provide the posterior probabilities of each functional class averaged across all the noncoding and synonymous coding variants in the training set. The predicted MACIE score for a given variant can be interpreted as the posterior probability of that variant belonging to (0,0), neither conserved nor regulatory classes; (1,0), the conserved but not the regulatory class; (0,1), the regulatory but not the conserved class; and (1,1), both conserved and regulatory classes. The four MACIE scores necessarily sum up to 1. A chi-squared test comparing observed and expected percentages under independence of evolutionary conservation and regulatory classes gives a significant *P* value of less than 2.2×10^−16^, suggesting that the two classes are correlated. Since the observed percentage of functional variants that belong to (1,1) is statistically significantly greater than the expected percentage under independence (3.15% > 1.96%), we find strong evidence of enrichment of regulatory activity in conserved regions. Additionally, the MACIE model for noncoding and synonymous coding variants estimates that 8.05% and 24.34% of the variants show evolutionarily conserved and regulatory functionality, respectively. This is consistent with the prediction from LINSIGHT and other previous studies that approximately 7% - 9% of noncoding sites are under evolutionary constraint^20,33^, as well as an estimated upper bound of 25% of the functional fraction within the human genome^34^.

### ClinVar pathogenic and benign variants

We first validated our methods on a testing set consisting of all variants recorded in the ClinVar database^28,29^. Variant effect predictor (VEP) information was extracted from GENCODE^35^ and used to separate nonsynonymous coding variants from noncoding and synonymous coding variants in ClinVar. The two MACIE models described above were then applied to the respective partitions. We combined the ClinVar categories “pathogenic” and “likely pathogenic” into a single pathogenic class and treated these variants as the putatively functional class. Similarly, we combined the ClinVar categories “benign” and “likely benign” into a single benign class and treated these variants as the putatively non-functional class. The remaining variants were categorized as having uncertain significance.

We first tested MACIE’s ability to distinguish pathogenic variants (*n* = 33,714) from their benign counterparts (*n* = 14,410) among ClinVar nonsynonymous variants through two approaches. First, we calculated two marginal MACIE scores: (1) MACIE-damaging protein function score (denoted by MACIE-protein) as the sum of the posterior probabilities of “damaging protein functional/not conserved” and “damaging protein functional/conserved”; (2) MACIE-conserved score as the sum of the posterior probabilities of “damaging protein functional/conserved” and “not damaging protein functional/conserved”. We also considered the posterior probability of either damaging protein functional or conserved (denoted by MACIE-anyclass) by summing the posterior probabilities corresponding to at least one functional class. This example illustrates the versatility of MACIE’s posterior probability outputs, which can be summed to form new probability measures with various informative interpretations depending on the specific needs of each analysis.

**Figure 2** provides the receiver operating characteristic (ROC) curves and area under the curves (AUC) for the three MACIE approaches and seven one-dimensional scores for ClinVar nonsynonymous variants. Of the methods considered, MACIE-damaging protein function score delivered the highest discrimination power (AUC = 0.93), followed by CADD (AUC = 0.91), EIGEN (AUC = 0.90), and MACIE-anyclass (AUC = 0.89). These four methods substantially outperformed the supervised DANN (AUC = 0.78), the supervised FATHMM-XF (AUC = 0.74), and the evolution-based fitCons (AUC = 0.54). Similar results were observed when distinguishing between pathogenic missense (as opposed to all nonsynonymous) variants (*n* = 21,409) from their benign counterparts (*n* = 14,035) in ClinVar (**Supplementary Figure 2**).

**Figure 2.**
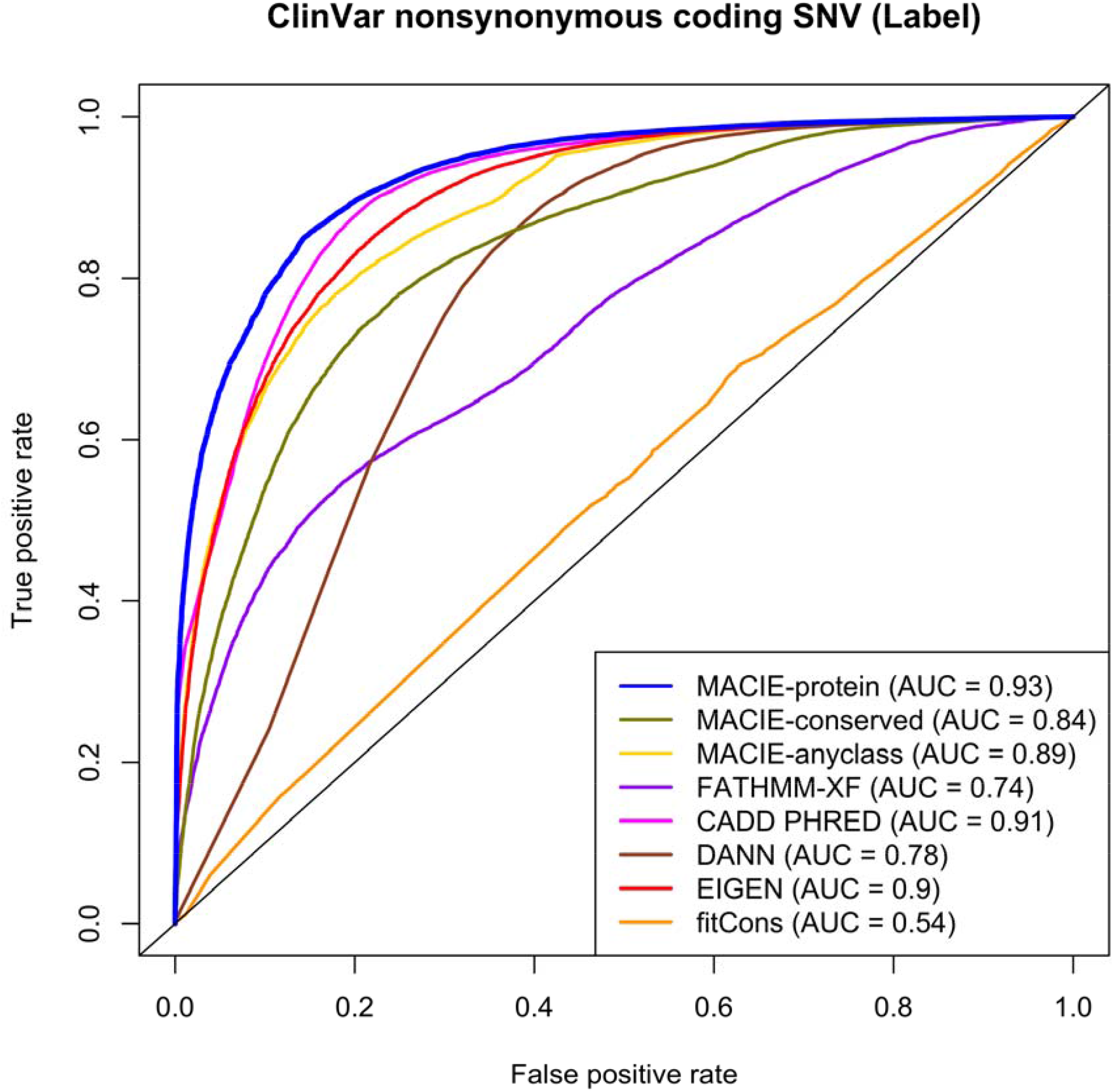
ROC curves comparing the performances of MACIE and other functional scores in discriminating between ClinVar pathogenic and benign nonsynonymous coding variants.

Next, we identified 40,109 noncoding variants from ClinVar database in total, including 6,551 pathogenic variants, and 33,558 benign variants. For these noncoding variants, we chose to calculate a marginal MACIE-conserved score, as ClinVar pathogenic noncoding variant labels track closely with evolutionary conservation scores (**Figure 1**). ROC curves and AUCs for discriminating between the pathogenic and benign variants are provided in **Supplementary Figure 3**. MACIE-conserved score showed comparable performance (AUC = 0.95) to FATHMM-XF score, which showed the highest discrimination power (AUC = 0.97). The outperformance of FATHMM-XF in this specific example should be expected because FATHMM-XF is a supervised machine-learning method trained on labels that bear many similarities to the labels defined in ClinVar, while MACIE is an unsupervised method. We performed Wilcoxon rank-sum tests to compare the distribution of integrative scores between ClinVar pathogenic and benign noncoding variants for each method. The Wilcoxon test *P* values for both FATHMM-XF and MACIE-conserved scores were less than 2.2 × 10^−308^, representing high discriminative abilities for each score. MACIE-conserved score substantially outperformed the unsupervised method EIGEN (AUC = 0.84) and the evolution-based method fitCons (AUC = 0.55).

### Loss-of-function nonsynonymous coding variants in *BRCA1*

We evaluated MACIE’s performance in predicting the deleteriousness of nonsynonymous coding variants located within 13 exons that encode functionally critical domains of *BRCA1*. A two-component Gaussian mixture model was fit based on the saturation genome editing function scores to classify all *BRCA1* variants as loss-of-function (LOF), intermediate (INT), or functional (FUNC), in a decreasing order of severity^30^. Thus, FUNC corresponds to benign variants in this experiment. We selected reported LOF nonsynonymous coding variants (*n* = 674) as the putative functional set and designated FUNC nonsynonymous coding variants (*n* = 1,443) as the putative non-functional set. Among all the methods compared (**Figure 3**), MACIE-damaging protein function score showed the highest predictive power (AUC = 0.91), followed by EIGEN (AUC = 0.88) and MACIE-anyclass (AUC = 0.88). The top three scores were much more powerful than CADD (AUC = 0.78), FATHMM-XF (AUC = 0.69), DANN (AUC = 0.60) and function score was the lowest (*P* = 7.60 × 10^−203^), and was orders of magnitude fitCons (AUC = 0.42). The Wilcoxon test *P* value for MACIE-damaging protein smaller than EIGEN (*P* = 7.22 × 10^−179^), CADD (*P* = 1.81 × 10^−95^) and other integrative scores. We observed similar results when distinguishing between *BRCA1* LOF nonsynonymous coding variants (*n* = 674) and ClinVar benign nonsynonymous coding variants (*n* = 14,410) (**Supplementary Figure 4**).

**Figure 3.**
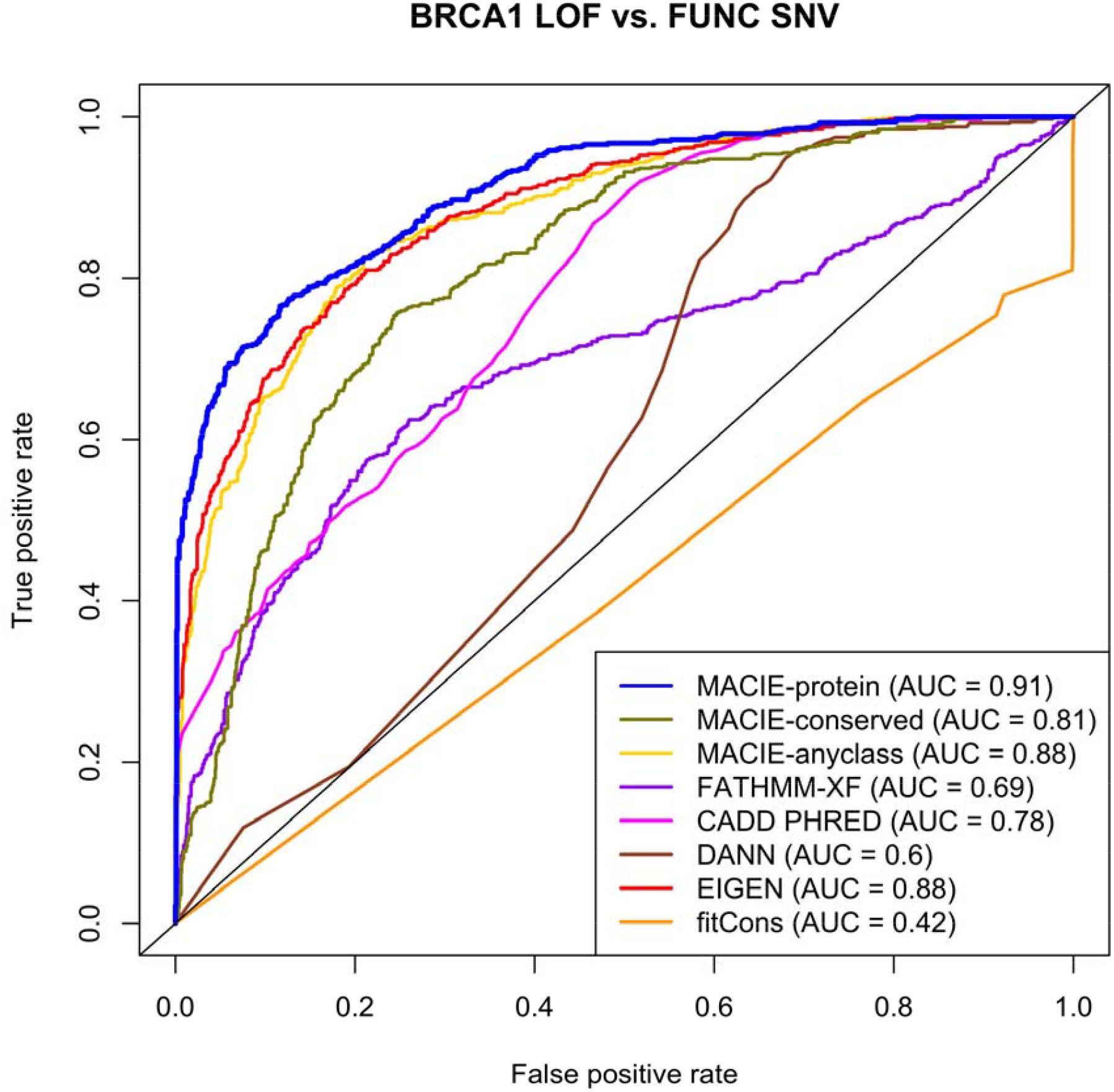
ROC curves comparing the performances of MACIE and other functional scores in discriminating between loss-of-function (LOF) and functional (FUNC) nonsynonymous coding variants within 13 exons that encode functionally critical domains of *BRCA1* based on saturation genome editing (SGE) data. Here the LOF class is our putative functional class and the FUNC class is our putative non-functional class.

### FANTOM5 CAGE-defined promoters and enhancers among 1000 Genomes noncoding variants

We tested the ability of MACIE to identify promoter regions defined by the cap analysis of gene expression conducted during the FANTOM5 project^7,8^. A total of 110,895 out of approximately 80 million noncoding variants from the 1000 Genomes Project Phase 3 data^36^ were mapped to such regions and therefore labeled as CAGE promoters. For each identified CAGE promoter variant, we used the 1000 Genomes Project database to randomly select a matched control variant (non-promoter) that possessed the same minor allele frequency (MAF) and same minimum distance to any gene transcription start site that was located at least 500 kilobase (kb) away from the promoter variant, yielding a total number of 97,298 variants in the control set (it was not possible to find a matched control for each CAGE variant). Similar to the previous analysis, we calculated a marginal MACIE-regulatory score by summing the two probabilities corresponding to the regulatory class (denoted by MACIE-regulatory). ROC curves and AUCs for discriminating between CAGE promoters and non-promoters are provided in **Figure 4a**. MACIE-regulatory and MACIE-anyclass scores showed the highest discrimination power (AUC = 0.75), followed by EIGEN with AUC = 0.74. The Wilcoxon test *P* value for MACIE-regulatory score was less than 2.2 × 10^−308^, indicating high discrimination ability. FATHMM-XF (AUC = 0.54) and fitCons (AUC = 0.56) scores performed poorly due to the inability of these one-dimensional scores to capture epigenetic functionality. We also performed a similar analysis by contrasting CAGE-identified enhancers(*n* = 520,987) versus non-enhancers (*n =* 448,253) using noncoding variants from the 1000 Genomes Project. The results were similar, with MACIE-regulatory score displaying the highest predictive power and significantly outperforming all other state-of-the-art methods (**Figure 4b**).

**Figure 4.**
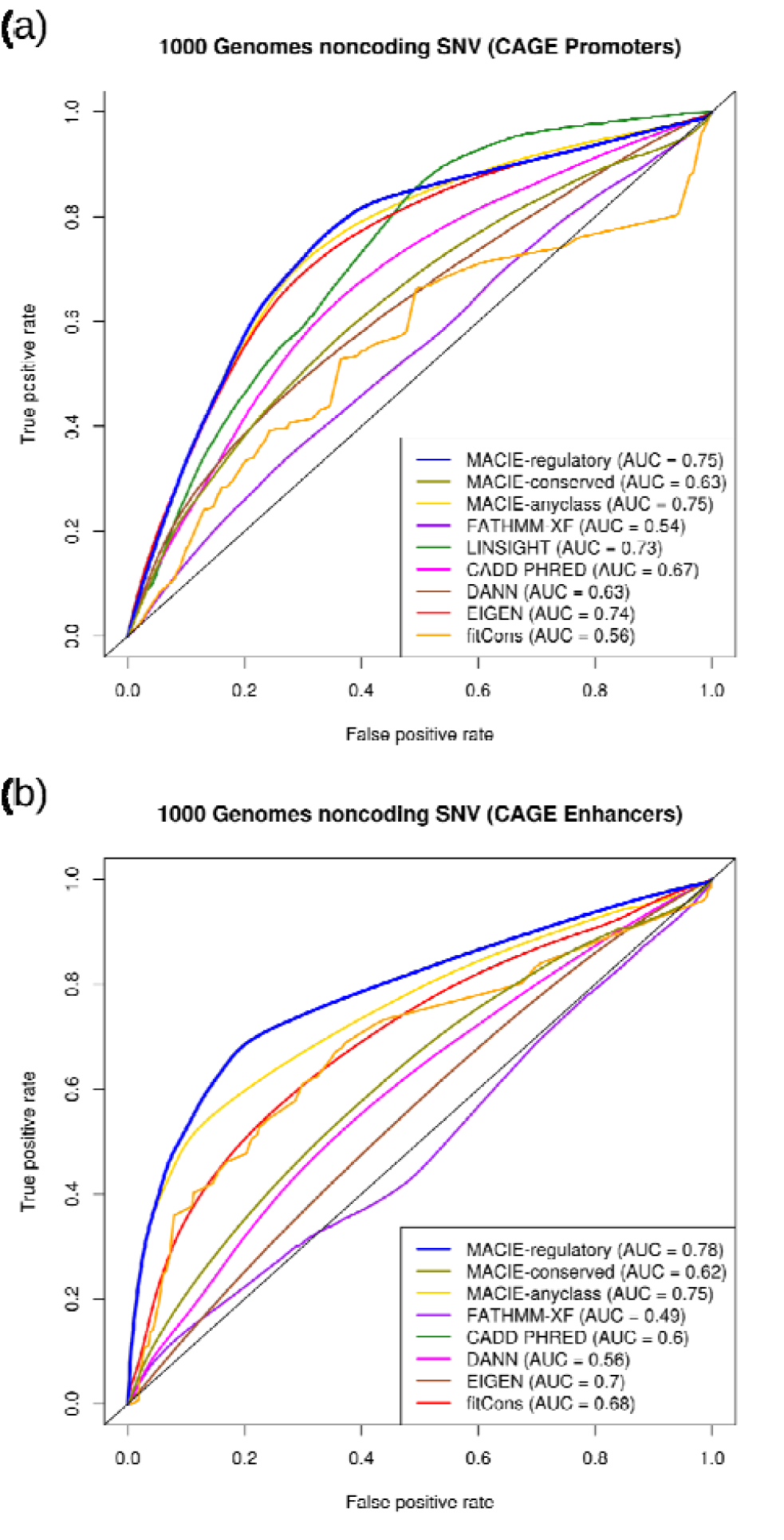
ROC curves comparing the performances of MACIE and other functional scores in discriminating between (a) CAGE identified promoters and non-promoters and (b) CAGE identified enhancers and non-enhancers among noncoding variants from 1000 Genomes Project Phase 3 data. For CAGE Enhancer predictions, LINSIGHT was excluded as it uses the FANTOM5 enhancer label as one of the genomic features in building the LINSIGHT score.

### MPRA validated variants and dsQTLs in lymphoblastoid cell lines

We examined the performance of MACIE for predicting cell type/tissue-specific regulatory variants using test sets from the massively parallel reporter assay. The MPRA dataset included validated regulatory variants in lymphoblastoid cell lines (LCLs)^31^. We paired each positive variant (*n* = 693) with four control variants from MPRA where neither allele showed significant differential expression at a Bonferroni corrected *P* value threshold of 0.1 (*n* = 2,772)^37^. **Figure 5a** shows that MACIE-regulatory score produced the highest discrimination power (AUC = 0.68), outperforming the second-best performing method (LINSIGHT, AUC = 0.64).

**Figure 5.**
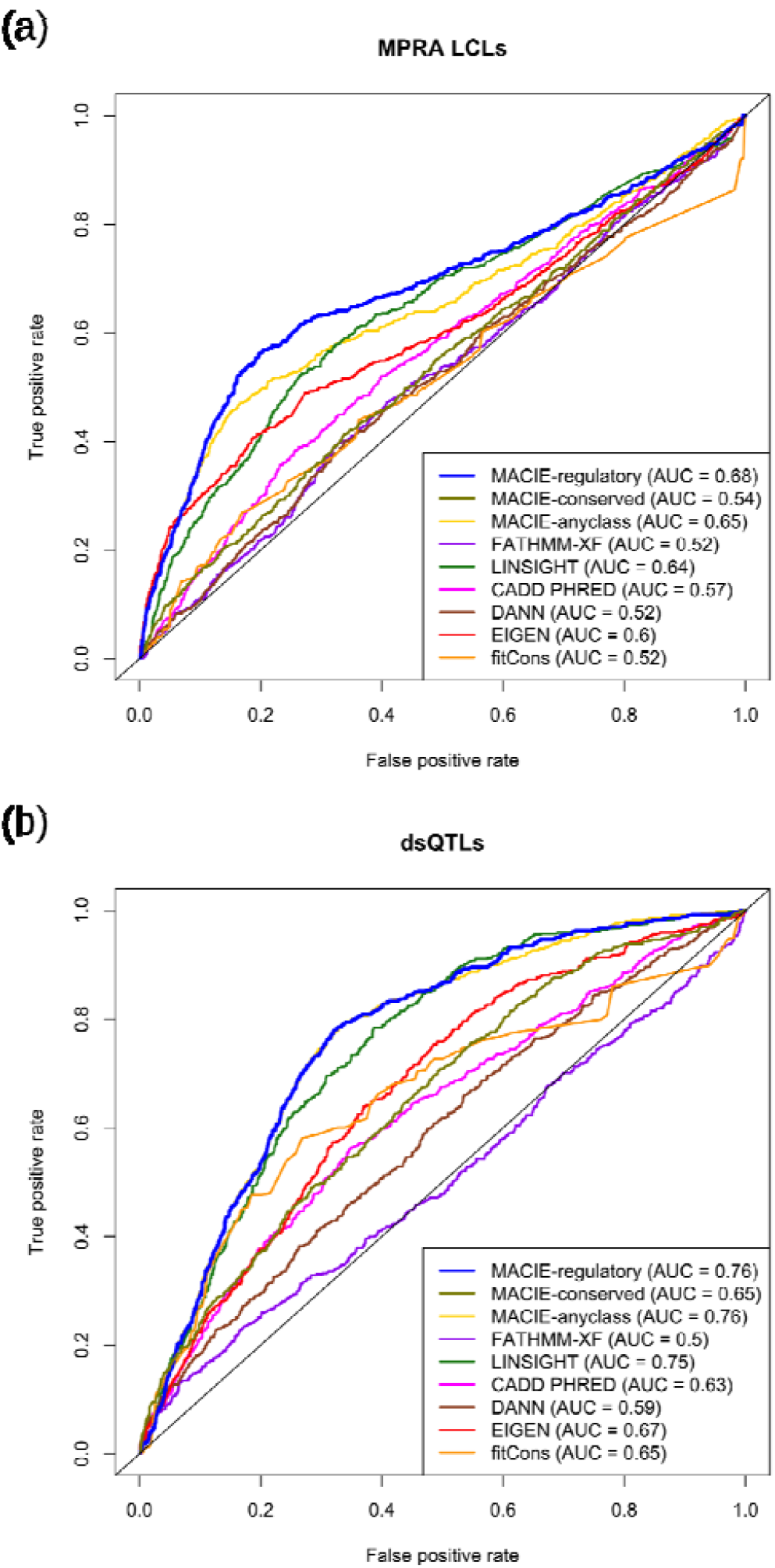
ROC curves comparing the performances of MACIE and other functional scores for prediction of (a) validated regulatory variants in lymphoblastoid cell lines (LCLs) from massively parallel reporter assays (MPRAs) and (b) dsQTLs identified using DNase I sequencing data in LCLs against control variants.

Finally, we evaluated the performance of our proposed method on a collection of dsQTLs that were identified using DNase I sequencing data from human lymphoblastoid cell lines^38^. Variants possessing association *P* values less than 1 × 10^−5^and residing within 100 bp of their corresponding DNase I-hypersensitive sites were chosen as the putatively functional set (*n* = 560)^39^. The control set of variants was randomly selected from a larger set of common variants (MAF > 5%) falling in the top 5% of DNase I sensitivity sites used to identify dsQTLs in the original study (*n* = 2,240). We observed that MACIE-regulatory score exhibited a larger AUC (AUC = 0.76) than all other methods (**Figure 5b**). MACIE-anyclass score also delivered robust performance on MPRA validated and dsQTLs datasets.

In summary, MACIE consistently ranked as one of the most powerful, robust and interpretable methods across a variety of settings and scientific questions. Our results show that while one-dimensional scores have gaps in coverage, a multi-dimensional scoring method offers robust and interpretable predictive performance. The ability of MACIE to interrogate variant functionality from multiple perspectives, at a level that is highly competitive with or better than state of the art methods, is unmatched by existing integrative functional scoring methods.

### MACIE prioritizes functional variants using lipids GWAS data

To illustrate the utility of MACIE scores in identifying plausible functional causal variants in genetic association studies, we applied MACIE to the publicly available lipids GWAS data from the European Network for Genetic and Genomic Epidemiology (ENGAGE) Consortium^40^. This dataset consists of lipids GWAS summary statistics for 9.6 million single nucleotide variants (SNVs) across 62,166 samples (**Supplementary Table 4**). We focused on genome-wide significant (*P*<5×10^−8^) SNVs associated with low-density lipoprotein cholesterol (LDL-C), high-density lipoprotein cholesterol (HDL-C), triglycerides (TG), and total cholesterol (TC). In total, we found 8, 9, 6, and 11 nonsynonymous coding SNVs that were predicted to belong to the protein damaging class with probability greater than 0.9 for LDL-C, HDL-C, TG, TC, respectively; 640, 377, 322, and 846 synonymous or noncoding SNVs that were predicted to belong to the regulatory class with probability greater than 0.9; 50, 64, 39, and 61 SNVs that were predicted to belong to the evolutionarily conserved class with probability greater than 0.9; and 9, 8, 10, 12 SNVs that were predicted to belong to both evolutionarily conserved and regulatory class with probability greater than 0.9 (**Supplementary Tables 5-12**). Compared to the total number of marginally significant SNVs for each trait (**Supplementary Table 4**), the MACIE scores reduce the number of SNVs prioritized for follow-up by an order of magnitude, saving much cost and effort in effectively pinpointing SNVs with relevant biological function.

For example, for LDL-C, the single most significant SNV was rs7412 (chr19:45412079 C/T; *P*<1×10^−316^). We predicted this known common missense SNV to be functional, as both MACIE-protein and MACIE-conserved scores provided a prediction greater than 0.95. These predictions highlight the multiple functional roles of this SNV. It is also worth noticing that the second most significant SNV rs1065853 (chr19:45413233 G/T; *P*<1×10^−316^) is in extremely high linkage disequilibrium (LD) with the leading SNV rs7412 (**Figure 6**). MACIE scores indicate that rs1065853 (upstream variant of *APOC1*) may possess a regulatory role since its MACIE-regulatory score is greater than 0.99, possibly suggesting that both the missense and regulatory variants can be putatively causal in affecting LDL-C levels. Similar results were observed for TC(**Supplementary Figure 5**). For HDL-C, although the single most significant SNV was rs17231506 (chr16:56994528 C/T; *P* = 6.88 × 10^−316^), the MACIE prediction was less than 0.01 for both classes. By scanning across the *CETP* locus and nearby noncoding regions associated with HDL-C, we found that two SNVs, rs72786786 (chr16:56985514 G/A; *P* = 2.52×10^−253^) and rs12720926 (chr16:56998918 A/G; *P* = 1.89 × 10^−260^), both under moderate to high LD with the leading SNV (**Supplementary Figure 6**), possess a MACIE-regulatory score greater than 0.99. These two SNVs may be more functionally important than rs17231506 and may provide more information regarding risk-perturbing biological mechanisms associated with this locus and can be prioritized for the leading SNV rs964184 (chr11:116648917 G/C; *P* = 1.74×10^−157^) in the functional follow-up. For TG, there is also a lack of functional evidence for the at this *APOA1/C3/A4/A5* gene cluster region. However, a SNV rs2075290(chr11:116653296 C/T; *P* = 2.13×10^−103^) in moderate LD with rs964184 at this locus has a MACIE-regulatory score of 0.88 (**Supplementary Figure 7**). These examples illustrate how MACIE scores can be used to supplement previous literature and provide additional information to aid prioritization of putatively functional causal variants for functional follow-up.

**Figure 6.**
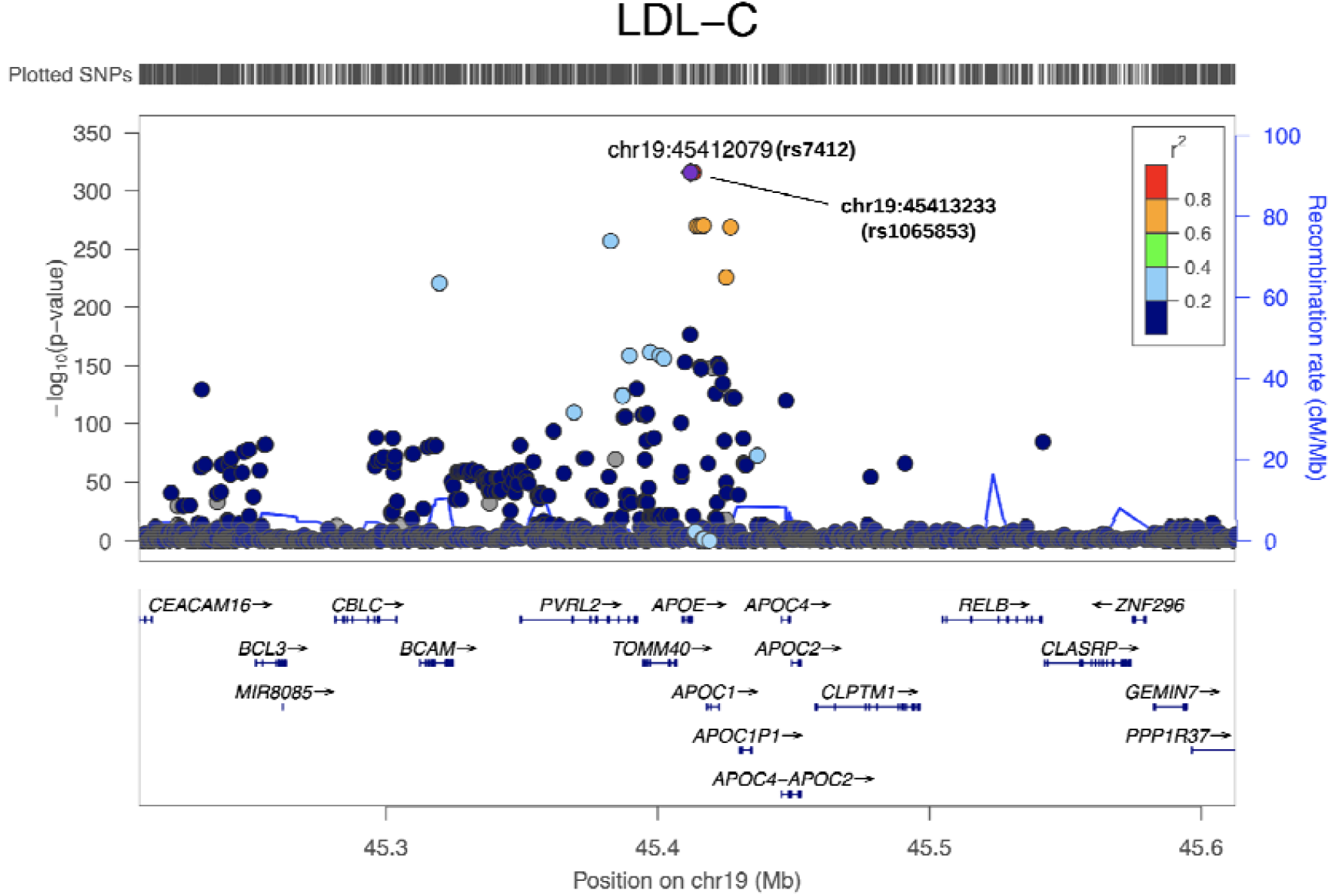
LocusZoom plot^49^ for GWAS associations of LDL-C at the *APOE* locus. The lipids GWAS summary statistics were from the European Network for Genetic and Genomic Epidemiology (ENGAGE) Consortium (*n* = 58,381)^40^. The MACIE-protein and MACIE-conserved scores for rs7412 are 0.96 and 0.97, respectively. The MACIE-conserved and MACIE-regulatory scores for rs1065853 are < 0.01 and > 0.99, respectively.

## Discussion

As the amount of publicly available annotation data increases and our understanding of variant functional effects continues to grow, describing variant functionality with a flexible yet practically interpretable and intuitive vocabulary will only become more important. Existing one-dimensional integrative scores cannot capture the multi-faceted functional profile of a variant because such ratings necessarily combine diverse, and possibly unrelated, sets of annotations into a single outcome. Oftentimes, they also ignore or do not fully take into account the correlation between individual annotations. Current supervised methods further demonstrate performance profiles that are linked strongly to the quality of training set labels. These supervised scores may lack robustness in the absence of gold-standard training sets.

In this paper we have proposed MACIE, an unsupervised multivariate mixed model framework that allows for multiple, possibly correlated, binary functional statuses. This framework offers several fundamental advancements over existing methods. First, MACIE provides multi-dimensional scores that measure functionality across multiple different functional classes. As posterior predictive probabilities, these scores are interpretable and scientifically relevant. They can be further summarized into marginal measures such as “probability of function according to at least one class of annotations” or “probability of function according to all classes of annotations”. Classes of annotation can be defined separately for different types of variants, for example, coding and noncoding variants.

Second, the MACIE model accommodates correlations both within- and between-classes. It has been reported that, while some of the available annotations measure similar notions of functionality, others provide distinct and complementary information^9,22^. By flexibly modeling potential, complex correlations across all the annotations, MACIE reflects this underlying biology. In doing so, it is better able to assign each annotation and group of annotations the appropriate amount of influence.

In multiple independent testing datasets, we showed that MACIE delivers powerful and robust performance in discriminating between functional and non-functional variants. Using lipids GWAS summary statistics data from the ENGAGE consortium, we also illustrated that MACIE offers an effective tool for fine-mapping studies to prioritize top hit in silico functional variants for experimental follow-up. MACIE scores have already been used, for example, to identify and characterize inflammation and immune-related risk variants in squamous cell lung cancer^41^. Finally, the proposed MACIE scores can be used as a weighting scheme to further empower variant-set analyses of rare variants^42^.

Our proposed MACIE framework provides a multi-dimensional functional class extension of several existing unsupervised single scoring frameworks, such as EIGEN^15^. MACIE fits a mixed model to the set of annotations for several latent functional classes and outputs the corresponding posterior component probabilities, which are highly interpretable. If we assume that there exists a single latent dichotomous variable summarizing functional status and that all annotations are independent conditional on the univariate functional status, then MACIE reduces to the GenoCanyon framework (**Supplementary Figure 1b**)^16^.

The versatility of the MACIE approach does introduce additional decisions that investigators need to make. For example, one needs to decide which set of annotations to include and how to group the annotations. The exponential family assumption in the model may also require a proper transformation for each individual annotation score before fitting the model. Operationally, users need to consider the trade-off between a more complex model (e.g., by increasing the number of classes or the number of functional scores in each class) and computation time. Such issues will become more relevant when extending the MACIE framework to integrate cell type-specific, tissue-specific, species-specific, or phenotype-related annotations^17,18,37^. Nevertheless, these choices again highlight the flexibility of the MACIE approach. Unlike other one-dimensional algorithms that rely on assumptions more likely to be satisfied when the number of annotations is small, the MACIE statistical model scales well with increasing annotation data. Thus, MACIE can be expected to provide more meaningful predictions as the availability of annotation scores continues to expand and the quality of these data improves.

A final important consideration in practical analysis concerns the differences between supervised and unsupervised methods. The performance of unsupervised scores may lag behind supervised methods when training datasets with relevant, high-quality labels are available. We observed this behavior when comparing MACIE to FATHMM-XF in ClinVar noncoding variants. Future extensions of interest include development of tools capable of integrating both supervised and unsupervised methods to further improve prediction accuracy^37^.

The MACIE scores used in all benchmarking examples are available for download at https://github.com/xihaoli/MACIE. Precomputed MACIE scores for all nonsynonymous coding, synonymous coding and noncoding variants in the human genome will be available. All genomic coordinates are given in NCBI Build 37/UCSC hg19.

### URLs

1000 Genomes: http://www.1000genomes.org. BRCA1 SGE: https://sge.gs.washington.edu/BRCA1. CADD: http://cadd.gs.washington.edu. ChromHMM: http://compbio.mit.edu/ChromHMM. ClinVar: https://www.ncbi.nlm.nih.gov/clinvar. DANN: https://cbcl.ics.uci.edu/public_data/DANN. EIGEN: http://www.columbia.edu/~ii2135/eigen.html. ENGAGE Consortium: http://diagram-consortium.org/2015_ENGAGE_1KG. FANTOM5 CAGE: https://fantom.gsc.riken.jp/5/data. FATHMM-XF: http://fathmm.biocompute.org.uk/fathmm-xf. fitCons: http://compgen.cshl.edu/fitCons. GENCODE: https://www.gencodegenes.org. LINSIGHT:https://github.com/CshlSiepelLab/LINSIGHT. LocusZoom: http://locuszoom.org. MACIE: https://github.com/xihaoli/MACIE.

## Acknowledgments

This work was supported by grants R35-CA197449, U19-CA203654, U01-HG009088, and R01-HL113338 (to X. Lin), MH095797 and MH106910 (to I.I.L.).

## Author contributions

X. Li, G.Y., H.Z., I.I.L. and X. Lin designed the experiments. X. Li, G.Y., H.Z., I.I.L., and X. Lin performed the experiments. X. Li, G.Y., H.Z., R.S., Z.L., Y.L., I.I.L., and X. Lin acquired, analyzed, or interpreted data. X. Li, G.Y., R.S., I.I.L., and X. Lin wrote the manuscript. All authors contributed to manuscript editing and approved the manuscript.

## Additional information

Supplementary information for this paper includes Supplementary Figures (7 figures) and Supplementary Tables (14 tables).

## Methods

### The MACIE generalized linear mixed model (GLMM)

Suppose there are *N* genetic variants in total and we are interested in *M* latent annotation classes, each containing *L*_*j*_ annotation scores. For example, the first class may consist of *L*_1_ =4 protein functional scores and the second class may consist of *L*_2_ = 8 evolutionary conservation scores. For genetic variant *i* and annotation class *j*, we denote the set of *L*_*j*_ annotations as 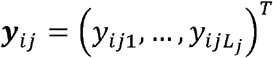 such that each variant is described by 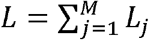 annotations in total. We want to estimate for each variant *i* the vector of binary functional statuses ***c***_*i*_ *= (c*_*i1*,_…*c*_*iM*_), where *c*_*ij*_ is the unobserved latent functional status for class *j*. Continuing our example, *c*_*i1*_ would denote membership in the evolutionarily conserved function class while *c*_*i2*_ would denote membership in the regulatory function class. Conditional on *c*_*ij*_ and a random effect variable *b*_*ijk*_, we assume that the elements of ***y***_*ij*_ are independent observations, each generated from a one-parameter exponential family with canonical parameterization. That is, for *j=1*,…,*M* and *k= 1*,…, *L*_*j*_

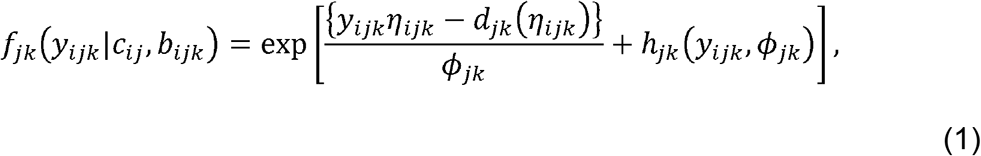

With

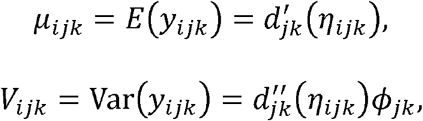

where *η*_*ijk*_ = *g*_*jk*_ (*μ*_*ijk*_) is a linear function of the functional status *c*_*ij*_ and random effect variable *b*_*ijk*_ such that

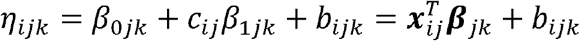

for ***x***_*ij*_ *= (1, c*_*ij*_)^*T*^ and ***β***_*jk*_ *= (β*_*0jk*_, *β*_*1jk*_)^*T*^. Additional correlations between elements of ***y***_*ij*_ are allowed by assuming that

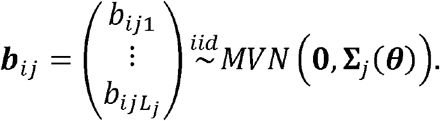

The marginal distribution of ***y***_*i*_ *=(****y***_*i1*_^*T*^,…,***y***_*iM*_^*T*^)^*T*^ can be obtained by integrating over the distribution of ***c***_*i*_ and ***b***_*i*_ *=(****b***_*i1*_^*T*^,…,***b***_*iM*_^*T*^)^*T*^,

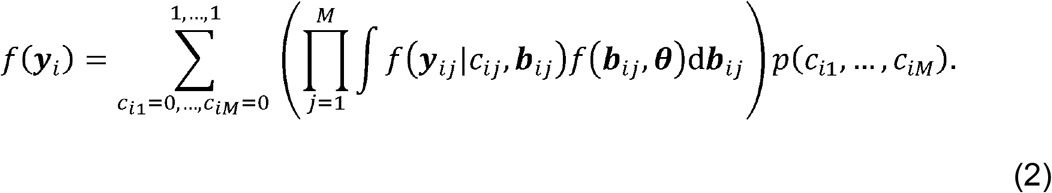

Our primary focus concerns calculation of *p(****c***_*i*_| ***y***_*i*_), the posterior probability of ***c***_*i*_ conditional on the observed data ***y***_*i*_. Because of the conditional independence of ***y***_*i*_ given ***c***_*i*_ and ***b***_*i*_ an Expectation-Maximization (EM) algorithm provides a natural approach^43^. However, the integration in Equation (2) cannot be evaluated in closed form whenever ***y***_*ij*_ conditional on *c*_*ij*_ and ***b***_*ij*_ is not normally distributed (e.g. ***y***_*ij*_ has dichotomous components). Thus, challenges arise in computing *p(****c***_*i*_|***y***_*i*_*)= f(****y***_*i*_|***c***_*i*_*)p(****c***_*i*_) / *f(****y***_*i*_). Approximations are used when applying the EM algorithm to obtain parameter estimates for non-normally distributed annotations.

Given the fitted model parameters and the full set of annotation scores for a new Variant *i’*, the MACIE score of variant *i’* is defined as the posterior probability vector *p*(***c***_***i***_***′ = z***|***y***_***i***_***′), z*** *∈ {0,1}*^*M*^. It can be calculated by performing one additional iteration of the EM algorithm.

### Derivation of the EM algorithm used in MACIE GLMM

In the following, we let **1**_*m*_ be the vector of length *m* where each element takes the value 1, and let **J**_*m*_ be the *m ×m* matrix of ones, i.e. 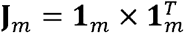. Let **I**_*m*_ be the *m ⨯m* identity matrix. Subscripts are dropped whenever the dimensions of the vector or matrix are clear. Our derivations follow those of Sammel et al.^44^, who considered a general class of latent variable models that allow for linear effects of covariates on multiple outcomes.

### Maximization step

If ***c***_*i*_ and ***b***_*ij*_ were directly observable, one can maximize the complete data log-likelihood,

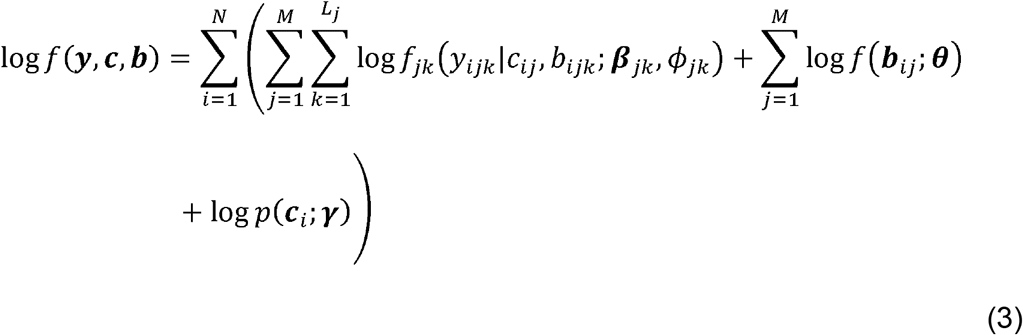

to estimate the unknown parameters ***ζ*=(*β***,***ϕ*** ,***γ***, ***θ*)** where *γ* corresponds to the vector of length 2^*M*^ that holds the probability of each possible realization of ***c***_*i*_ However, since ***c***_*i*_ and ***b***_*ij*_ are unobservable, the EM algorithm can be applied by instead solving the expected score functions, where the expectation is taken with respect to

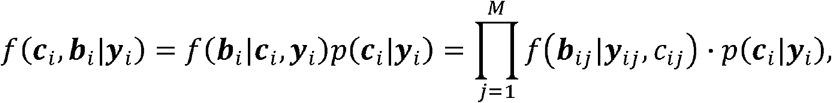

which is the posterior distribution of the missing data conditional on the observed data^45^. If we let *S*_*i*_(***ζ***) denote the complete data score function ∂ log *f(****y***_*i*_, ***c***_*i*_, ***b***_*i*_) */* ∂***ζ*** of Equation (3) for the *i*th variant, then each variant’s contribution to the expected score function for ***γ*** is given by

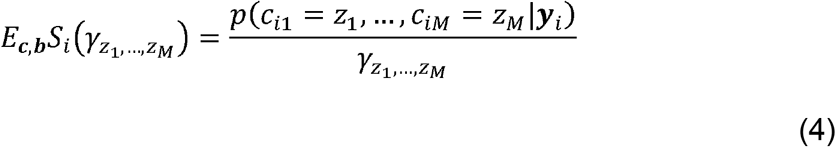

for all *(z*_*1*_,…*z*_*M*_) ∈ {0,1}^*M*^ .Therefore, based on Equation (4), the M-step update for ***γ*** in moving from iteration *r* to *r*+*1* is

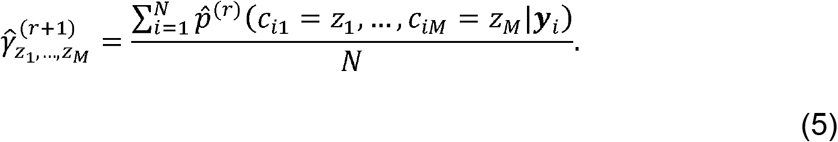

For ***ζ***_*jk*,_ the subset of parameters corresponding to only the *jk*th outcome, the contribution to the expected score equation for variant *i* is

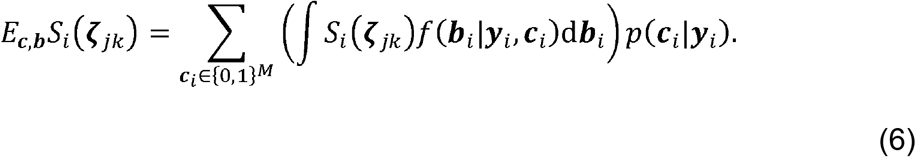

Depending on the form of the score function associated with the complete data log-likelihood 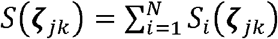, the solution to *E*_*c,b*_*S*(***ζ***_*jk*,_)= **0** may or may not be available in closed form. In the absence of a closed form solution, we update the estimates ***ζ***_*jk*,_ through a one-step Fisher scoring algorithm. The usual method of estimation for this model is iteratively reweighed least squares,^46^ where the weight function is updated at every iteration.

### Expectation step

Given the current estimates of the parameters, ***ζ=(β,ϕ ,γ, θ***) the E-step is complicated by the need to compute expectations with respect to the posterior distributions *f(****b***_*i*_|***y***_*i*_,***c***_*i*_) and *p(****c***_i_|***y***_*i*_) of the missing data, conditional on the observed data. Only for normal outcomes will the posterior distributions have closed form solutions. In our setting, there are generally no closed form expressions for *f(****b***_*i*_|***y***_*i*_,***c***_*i*_) and *p(****c***_*i*_|***y***_*i*_) As an alternative to analytical solutions, we first write the expectation of functions of the data *g*(***c***_*i*_,*b*_*i*_*)= g (****y***_*i*_,***c***_*i*_,***b***_*i*_) by,

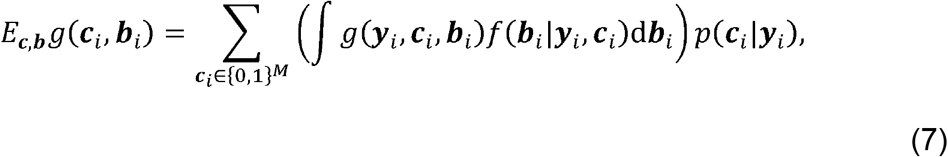

and further rewrite the posterior distributions as

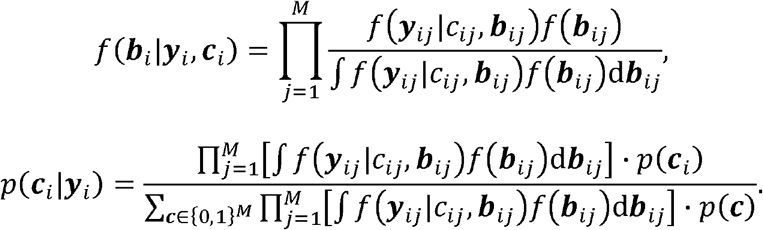

By substituting into Equation (7), we obtain

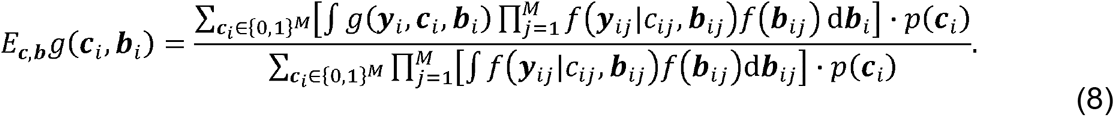

If g (***y***_*i*_,***c***_*i*_,***b***_*i*_) = g (***y***_*ij*′_,***c***_*ij*′_,***b***_*ij*′_) for some *j*’ ∈ {1, …,*M*}, then the integral in the numerator of Equation (8) is equivalent to

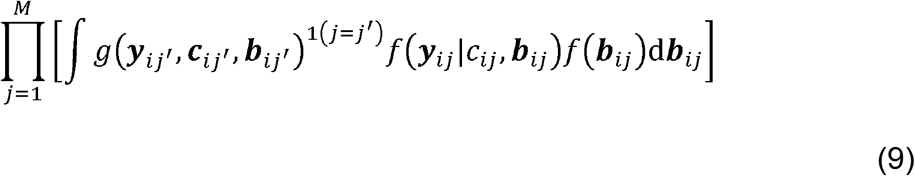

where 1*(j =j’)* is equal to 1 if *j =j’* and 0 otherwise. In this case, a practical approximation is to use multivariate Gauss-Hermite quadrature. To approximate Equation (9), we select *T* fixed abscissae 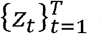 and corresponding approach for weights 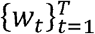 for a quadrature whose integration kernel is given by the density of a standard normal distribution^47^. Given the spectral decomposition of 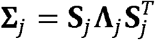 let *σ*_*jt*_ *= {σ*_*jt*_ *(1)*, …, *σ*_*jt*_ *(L*_*j*_*)}* be an ordered set of *L*_*j*_ integers obtained by sampling with replacement from *{1*,…,*T}*, 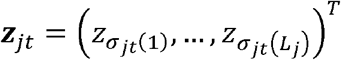 the corresponding set of abscissae, and 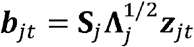. Then each term in the product of Equation (9)

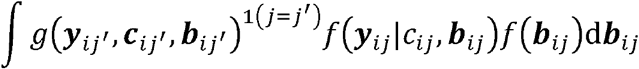

can be approximated as

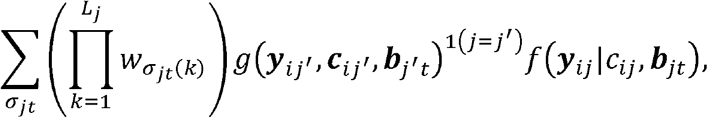

where the sum is over all the possible ordered sets *σ*_*jt*_. For some ordered sets *σ*_*jt*_ the weights 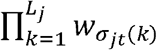 are very small and thus contribute little to the sum. We may choose to remove these quantities by pruning a specified fraction of the smallest weights.

### MACIE: EM algorithm for mixed binary and normal annotations

The general formulation of Equation (1) allows different link functions *g*_*jk*_(.) for different annotations, as well as different covariance structures **Σ**_*j*_ ***(θ)*** to accommodate for correlations between the annotations (**Supplementary Figure 1c**). In this section, we derive specific theoretical results for the EM algorithm when annotations are either conditionally bernoulli or normal random variables, i.e. all link functions *g*_*jk*_(.) are either the identity or logistic link. We also introduce restrictions on the covariance matrices **Σ**_*j*_ (***θ***) that allow for accurate approximations while greatly reducing the algorithm’s computational speed. We call this algorithm MACIE for Multi-dimensional Annotation Class Integrative Estimation.

Suppose that conditionally on *c*_*ij*_ and ***b***_*ij*_ the first 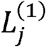 of the *L*_*j*_ outcomes ***y***_*i,j*_ follow a bernoulli distribution and the remaining 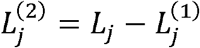 outcomes follow a normal distribution. That is, for 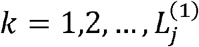, *y*_*ijk*_ has distribution

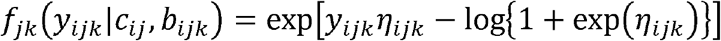

where *μ* _ijk_ = exp (*η*_*ijk*_) / {1 + exp (*η* _*ijk*_)} and *V*_*ijk*_ = *μ*_*ijk*_ (1−*μ*_*ijk*_). Then for 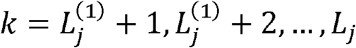, *y*_*ijk*_ has the distribution

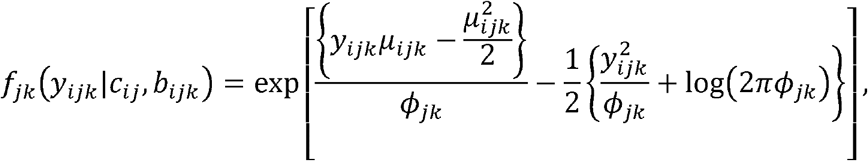

where *μ*_*ijk*_ = *η*_*ijk*_ and *V*_*ijk*_ = *ϕ*_*jk*._

If ***Σ***_*j*_ ***(θ)*** is left unstructured, then the EM algorithm will need to estimate *L*_*j*_ *(L*_*j*_ + *1)/2* parameters for the covariance matrix of class *j*. An even greater computational challenge is that the multivariate Gauss-Hermite quadrature will require 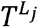 fixed abscissas. Thus, to reduce the number of model parameters and to make the algorithm computationally feasible, we assume that ***b***_*ij*_ = **Λ***_j_****f***_*ij*_ where ***f***_*ij*_ is an unobserved vector of length *P*_*j*_ *< L*_*j*_ that follows *MVN(0,1)*. Then for the E-step,

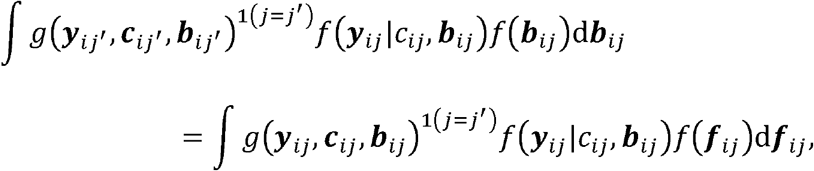

so that integration is over a *P*_*j*_-dimensional space as opposed to an *L*_*j*_-dimensional space. The assumption ***b***_*ij*_ = **Λ**_*j*_***f***_*ij*_ forms the basis of factor analysis models^48^ and is appropriate when the relationship between *L*_*j*_ manifest variables is thought to be primarily a result of the relationship between *P*_*j*_ underlying variables. For functional annotations, the underlying variables are likely to correspond to different approaches measuring the same element. As in factor analysis, the larger the factor loading *λ*_*jkp*,_the more the *jk*th annotation is said to “load” on the *p*th factor.

For the *L*_*j1*_ binary outcomes, substituting the appropriate quantities into Equation (6) leads to the following expected score functions for variant on outcome *jk* :

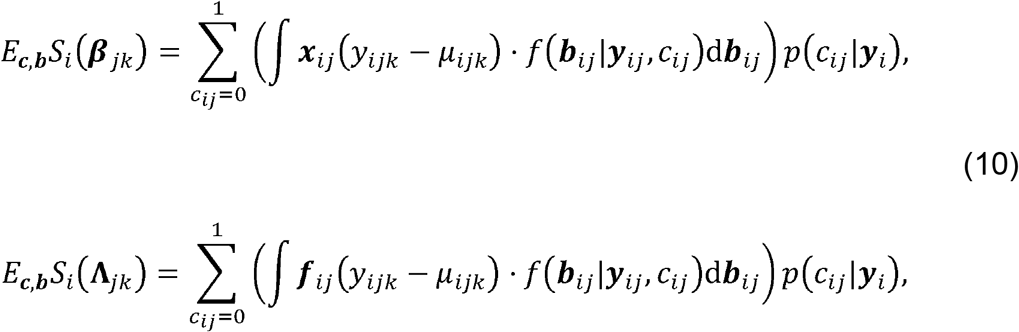

where **Λ**_*jk*_ is the *k*th column vector of 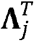.

To update estimates for ***β***_*jk*_ using a one-step Fisher scoring algorithm, we consider a Taylor series expansion of the expected score function (Equation (10)) about the true parameter ***β***_*jk*_

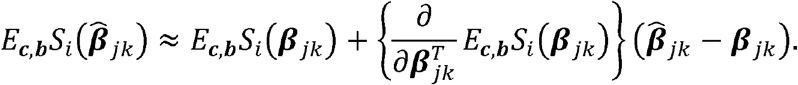

Since 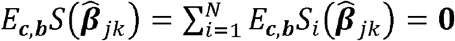, and assuming regularity conditions that allow the interchange of differentiation and integration, we have

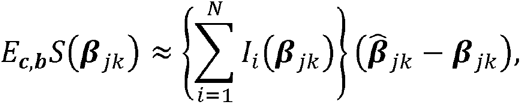

where *I*_*i*_ is the *i*th variant’s contribution to the observed data Fisher information associated with the *jk*th outcome:

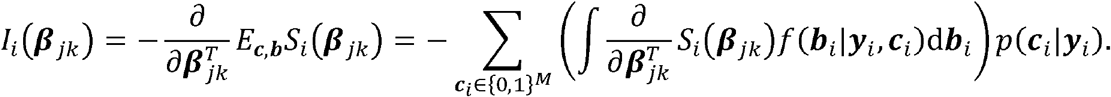

The expected information is obtained by taking an additional expectation with respect to the observed outcomes ***y***_*i*_ :

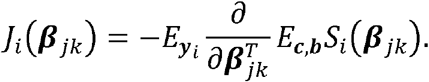

Interchanging derivatives and expectations yields

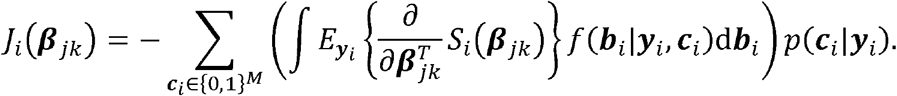

For binary outcomes with logistic link, the expected information is

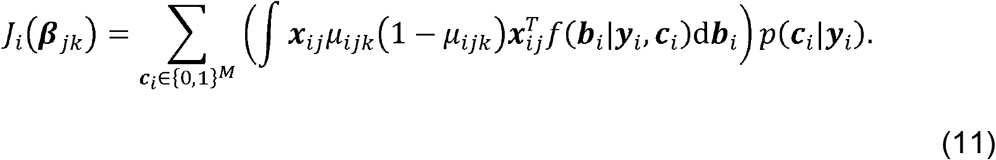

Equations (10) and (11) yield the following scoring algorithm at iteration *r*+*1:*

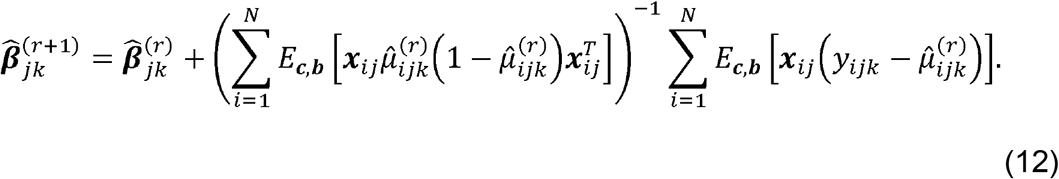

Similarly,

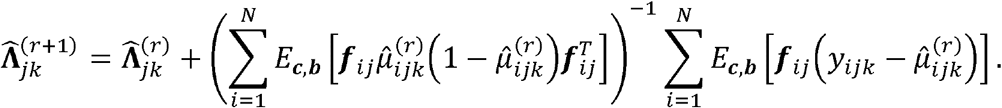

For the 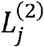 normal outcomes, contributions to the complete data score functions for each variant *i* are

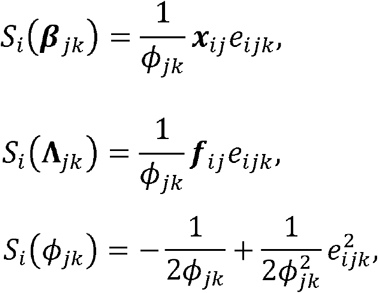

where 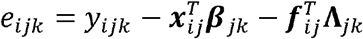, It follows that

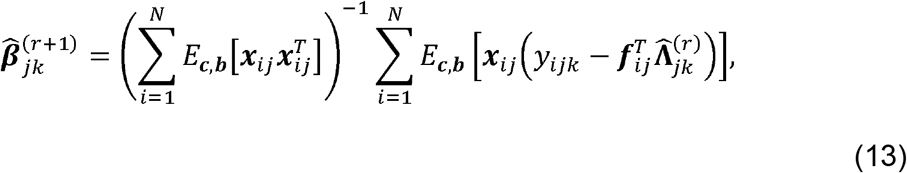

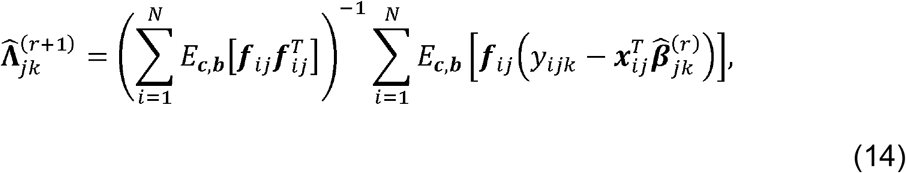

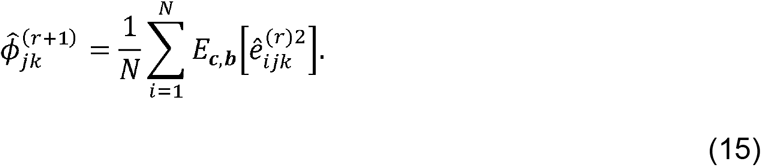

Beginning with reasonable initial estimates of the parameters, MACIE proceeds by first using the E-step to obtain the desired expectations relative to the posterior distribution. Given those estimates, MACIE then solves the expected score equations to obtain new parameter estimates or one-step updates a according to Equations (5), (13)-(15). The algorithm proceeds until the relative change in the estimated parameters is sufficiently small (<10^−4^) with a maximum of 200 iterations.

### Data analysis using the MACIE GLMM

We used the proposed framework to fit the MACIE GLMM models for (1) nonsynonymous coding variants and (2) noncoding and synonymous coding variants separately. For nonsynonymous coding variants, we considered fitting a two-class MACIE model *(M = 2)* where the damaging protein function class included four protein substitution scores: SIFT, PolyPhenDiv, PolyPhenVar (dichotomous) and Mutation Assessor (continuous), with two latent factors of ∑_1_; and the evolutionary conserved class included eight conservation scores: GERP_NR, GERP_RS, PhyloPri, PhyloPla, PhyloVer (continuous), and PhastPri,PhastPla, PhastVer (dichotomous), with two latent factors of **∑**_2_ (**Supplementary Table 1**). As such, the MACIE score predicted for each nonsynonymous coding variant is a vector of length 4, representing the estimated joint posterior probabilities of belonging to (0,1) - “not damaging protein functional and conserved”; (1,0) - “damaging protein functional and not conserved”; (0,0) - “not damaging protein functional and not conserved”; (1,1) - “both damaging protein functional and conserved”. The MACIE GLMM regression parameter estimates from the training set of nonsynonymous coding variants are presented in **Supplementary Table 13**.

For noncoding and synonymous coding variants, we considered fitting a two-class MACIE model *(M = 2)*, where the evolutionary conserved class included the same eight conservation scorers as the nonsynonymous coding model, with two latent factors of **∑**_1_, and the regulatory class included a total of twenty-eight transformed (continuous) epigenetic scores scores, consisting of three histone marks and 12 open chromatin marks from the ENCODE Project, three transcription factor binding site scores, GC content, CpG content, five chromatin state probabilities derived from the 15 state ChromHMM model, a background selection score, and physical distance metrics, with three latent factors of **∑**_2_ (**Supplementary Table 1**). As such, the MACIE score predicted for each noncoding or synonymous coding variant is also a vector of length 4, representing the estimated joint posterior probabilities of belonging to (0,1) - “not conserved and regulartory functional”; (1,0) - “conserved and not regulatory functional”; (0,0) - “not conserved and not regulatory functional”; (1,1) - “both conserved and regulatory functional”. The MACIE GLMM regression parameter estimates from the training set of noncoding and synonymous coding variants are presented in **Supplementary Table 14**.

## References

1. Adzhubei, I.A. et al. A method and server for predicting damaging missense mutations. Nature methods 7, 248 (2010).

2. Siepel, A. et al. Evolutionarily conserved elements in vertebrate, insect, worm, and yeast genomes. Genome research 15, 1034–1050 (2005).

3. Pollard, K.S., Hubisz, M.J., Rosenbloom, K.R. & Siepel, A. Detection of nonneutral substitution rates on mammalian phylogenies. Genome research 20, 110–121 (2010).

4. Davydov, E.V. et al. Identifying a high fraction of the human genome to be under selective constraint using GERP++. PLoS computational biology 6, e1001025 (2010).

5. ENCODE Project Consortium. An integrated encyclopedia of DNA elements in the human genome. Nature 489, 57 (2012).

6. Kundaje, A. et al. Integrative analysis of 111 reference human epigenomes. Nature 518, 317 (2015).

7. Forrest, A.R. et al. A promoter-level mammalian expression atlas. Nature 507, 462 (2014).

8. Andersson, R. et al. An atlas of active enhancers across human cell types and tissues. Nature 507, 455 (2014).

9. Kellis, M. et al. Defining functional DNA elements in the human genome. Proceedings of the National Academy of Sciences 111, 6131–6138 (2014).

10. Kircher, M. et al. A general framework for estimating the relative pathogenicity of human genetic variants. Nature genetics 46, 310 (2014).

11. Quang, D., Chen, Y. & Xie, X. DANN: a deep learning approach for annotating the pathogenicity of genetic variants. Bioinformatics 31, 761– 763 (2014).

12. Ritchie, G.R., Dunham, I., Zeggini, E. & Flicek, P. Functional annotation of noncoding sequence variants. Nature methods 11, 294 (2014).

13. Shihab, H.A. et al. An integrative approach to predicting the functional effects of non-coding and coding sequence variation. Bioinformatics 31, 1536–1543 (2015).

14. Rogers, M.F. et al. FATHMM-XF: accurate prediction of pathogenic point mutations via extended features. Bioinformatics (2017).

15. Ionita-Laza, I., McCallum, K., Xu, B. & Buxbaum, J.D. A spectral approach integrating functional genomic annotations for coding and noncoding variants. Nature genetics 48, 214 (2016).

16. Lu, Q. et al. A statistical framework to predict functional non-coding regions in the human genome through integrated analysis of annotation data. Scientific reports 5, 10576 (2015).

17. Bodea, C.A. et al. PINES: phenotype-informed tissue weighting improves prediction of pathogenic noncoding variants. Genome Biology 19, 173 (2018).

18. Backenroth, D. et al. FUN-LDA: A latent Dirichlet allocation model for predicting tissue-specific functional effects of noncoding variation: methods and applications. The American Journal of Human Genetics 102, 920–942 (2018).

19. Gulko, B., Hubisz, M.J., Gronau, I. & Siepel, A. A method for calculating probabilities of fitness consequences for point mutations across the human genome. Nature genetics 47, 276 (2015).

20. Huang, Y.-F., Gulko, B. & Siepel, A. Fast, scalable prediction of deleterious noncoding variants from functional and population genomic data. Nature genetics 49, 618 (2017).

21. Tang, H. & Thomas, P.D. Tools for predicting the functional impact of nonsynonymous genetic variation. Genetics 203, 635–647 (2016).

22. Lee, P.H. et al. Principles and methods of in-silico prioritization of non-coding regulatory variants. Human genetics 137, 15–30 (2018).

23. Liu, X., Wu, C., Li, C. & Boerwinkle, E. dbNSFP v3. 0: A one-stop database of functional predictions and annotations for human nonsynonymous and splice-site SNVs. Human mutation 37, 235–241 (2016).

24. Ng, P.C. & Henikoff, S. SIFT: Predicting amino acid changes that affect protein function. Nucleic acids research 31, 3812–3814 (2003).

25. Reva, B., Antipin, Y. & Sander, C. Predicting the functional impact of protein mutations: application to cancer genomics. Nucleic acids research 39, e118–e118 (2011).

26. Ernst, J. & Kellis, M. Discovery and characterization of chromatin states for systematic annotation of the human genome. Nature biotechnology 28, 817 (2010).

27. McVicker, G., Gordon, D., Davis, C. & Green, P. Widespread genomic signatures of natural selection in hominid evolution. PLoS genetics 5, e1000471 (2009).

28. Landrum, M.J. et al. ClinVar: public archive of relationships among sequence variation and human phenotype. Nucleic acids research 42, D980–D985 (2013).

29. Landrum, M.J. et al. ClinVar: public archive of interpretations of clinically relevant variants. Nucleic acids research 44, D862–D868 (2015).

30. Findlay, G.M. et al. Accurate classification of BRCA1 variants with saturation genome editing. Nature 562, 217 (2018).

31. Tewhey, R. et al. Direct identification of hundreds of expression-modulating variants using a multiplexed reporter assay. Cell 165, 1519– 1529 (2016).

32. Kheradpour, P. et al. Systematic dissection of regulatory motifs in 2,000 predicted human enhancers using a massively parallel reporter assay. Genome research, gr. 144899.112 (2013).

33. Rands, C.M., Meader, S., Ponting, C.P. & Lunter, G. 8.2% of the human genome is constrained: variation in rates of turnover across functional element classes in the human lineage. PLoS genetics 10, e1004525 (2014).

34. Graur, D. An upper limit on the functional fraction of the human genome. Genome biology and evolution 9, 1880–1885 (2017).

35. Harrow, J. et al. GENCODE: the reference human genome annotation for The ENCODE Project. Genome research 22, 1760–1774 (2012).

36. Genomes Project Consortium. An integrated map of genetic variation from 1,092 human genomes. Nature 491, 56–65 (2012).

37. He, Z., Liu, L., Wang, K. & Ionita-Laza, I. A semi-supervised approach for predicting cell-type specific functional consequences of non-coding variation using MPRAs. Nature communications 9, 5199 (2018).

38. Degner, J.F. et al. DNase I sensitivity QTLs are a major determinant of human expression variation. Nature 482, 390 (2012).

39. Li, M.J. et al. Predicting regulatory variants with composite statistic. Bioinformatics 32, 2729–2736 (2016).

40. Surakka, I. et al. The impact of low-frequency and rare variants on lipid levels. Nature genetics 47, 589–597 (2015).

41. Sun, R. et al. Integration of multiomic annotation data to prioritize and characterize inflammation and immune-related risk variants in squamous cell lung cancer. Genetic Epidemiology, 1–16 (2020).

42. Li, X. et al. Dynamic incorporation of multiple in silico functional annotations empowers rare variant association analysis of large whole-genome sequencing studies at scale. Nature genetics 52, 969–983 (2020).

## References

43. Dempster, A.P., Laird, N.M. & Rubin, D.B. Maximum likelihood from incomplete data via the EM algorithm. Journal of the royal statistical society. Series B (methodological), 1–38 (1977).

44. Sammel, M.D., Ryan, L.M. & Legler, J.M. Latent variable models for mixed discrete and continuous outcomes. Journal of the Royal Statistical Society: Series B (Statistical Methodology) 59, 667–678 (1997).

45. Little, R.J. & Rubin, D.B. Statistical analysis with missing data. New York: Wiley, 1987 (1987).

46. McCullagh, P. & Nelder, J.A. Generalized Linear Models, Second Edition, (Taylor & Francis, 1989).

47. Abramowitz, M. & Stegun, I.A. Handbook of mathematical functions: with formulas, graphs, and mathematical tables, (Courier Corporation, 1964).

48. Lawley, D. & Maxwell, A. Factor analysis as a statistical method. Journal of the Royal Statistical Society. Series D (The Statistician) 12, 209–229 (1962).

49. Pruim, R.J. et al. LocusZoom: regional visualization of genome-wide association scan results. Bioinformatics 26, 2336–2337 (2010).

