## Supplementary Figures for "A multi-dimensional integrative scoring framework for predicting functional variants in the human genome"

**Supplementary Figure 1.** Models representing potential causal relations among annotations. **(a)** All annotations $\boldsymbol{y}_{ij}$ (e.g., conservation measures, epigenetic measures) are treated as consequences of a single latent dichotomous variable of function $\boldsymbol{c}_{i}$. Annotations are assumed to be independent conditional on $\boldsymbol{c}_{i}$, as proposed in GenoCanyon. **(b)** All annotations $\boldsymbol{y}_{ij}$ are treated as consequences of $\boldsymbol{c}_{i}$. Annotations may be correlated conditional on $\boldsymbol{c}_{i}$. **(c)** There are multiple, possibly related, latent dichotomous variables of function $c_{i1},\ldots,c_{iM}$. For each functional status $c_{ij}$, a subset of annotations $y_{ij1},\ldots,y_{ijL_{j}}$ are observed as consequences. Annotations measuring the same $c_{ij}$ may be correlated conditional on $c_{ij}$, as proposed in MACIE.

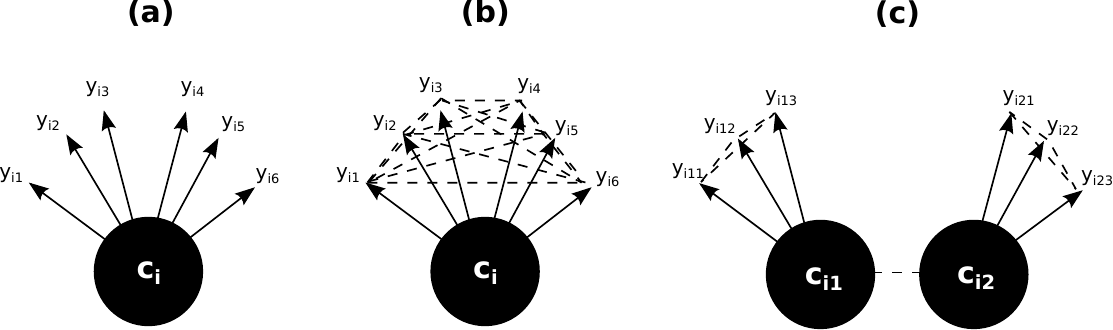

**Supplementary Figure 2.** ROC curves comparing the performances of MACIE and other functional scores in discriminating between ClinVar pathogenic and benign missense variants.

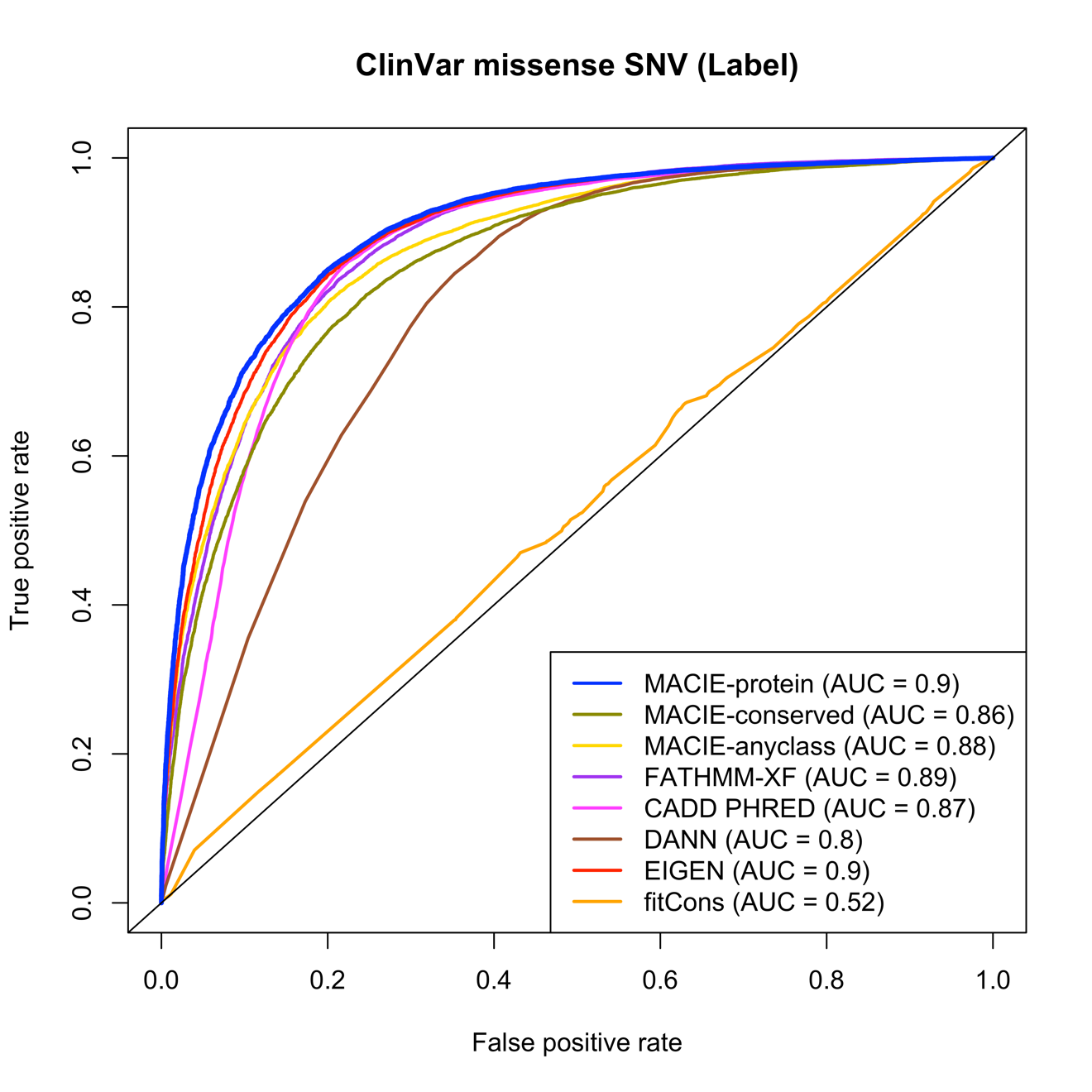

**Supplementary Figure 3.** ROC curves comparing the performances of MACIE and other functional scores in discriminating between ClinVar pathogenic and benign noncoding variants.

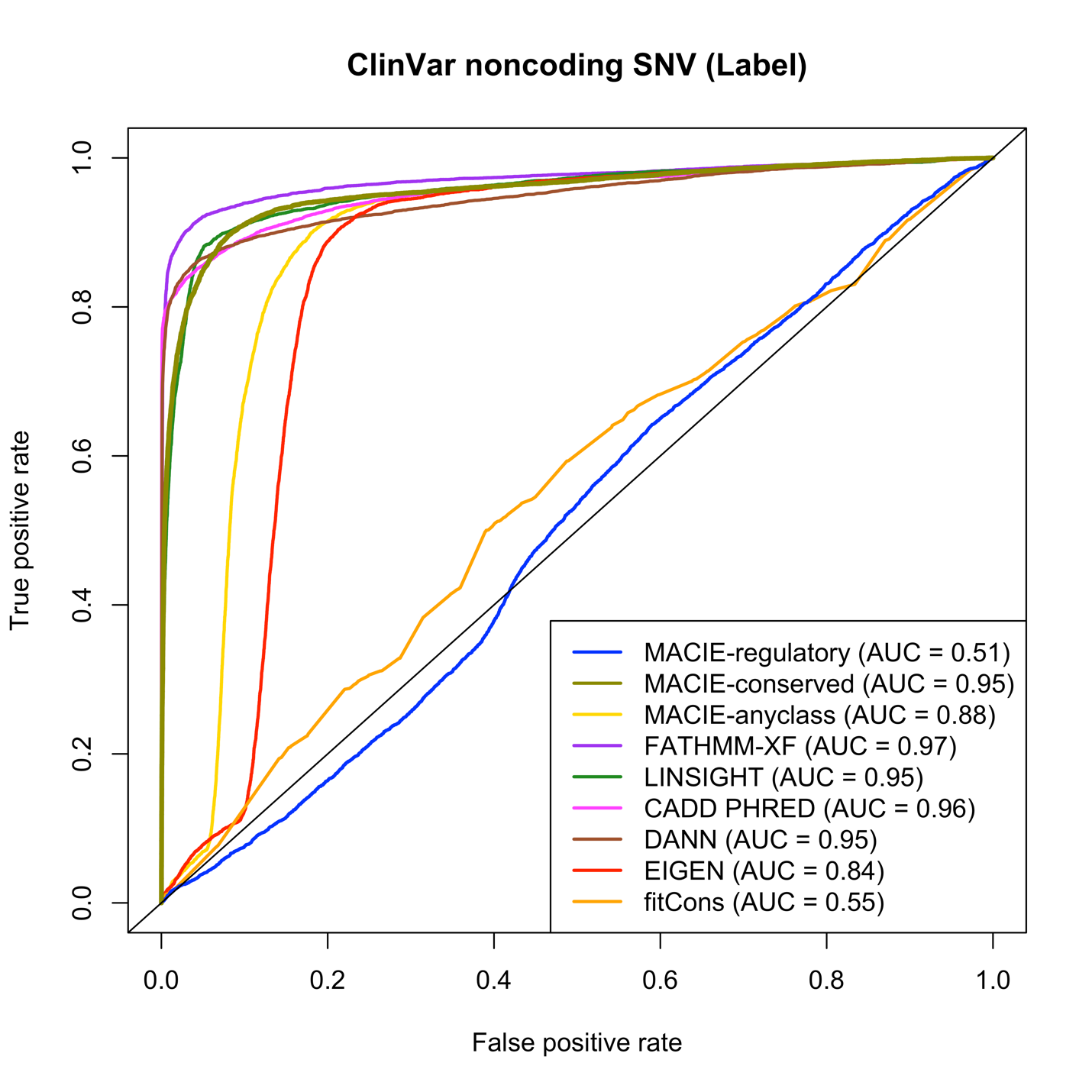

**Supplementary Figure 4.** ROC curves comparing the performances of MACIE and other functional scores in discriminating between loss-of-function (LOF) nonsynonymous coding variants within 13 exons that encode functionally critical domains of *BRCA1* (putative functional class) based on saturation genome editing (SGE) data and ClinVar benign nonsynonymous coding variants (putative non-functional class).

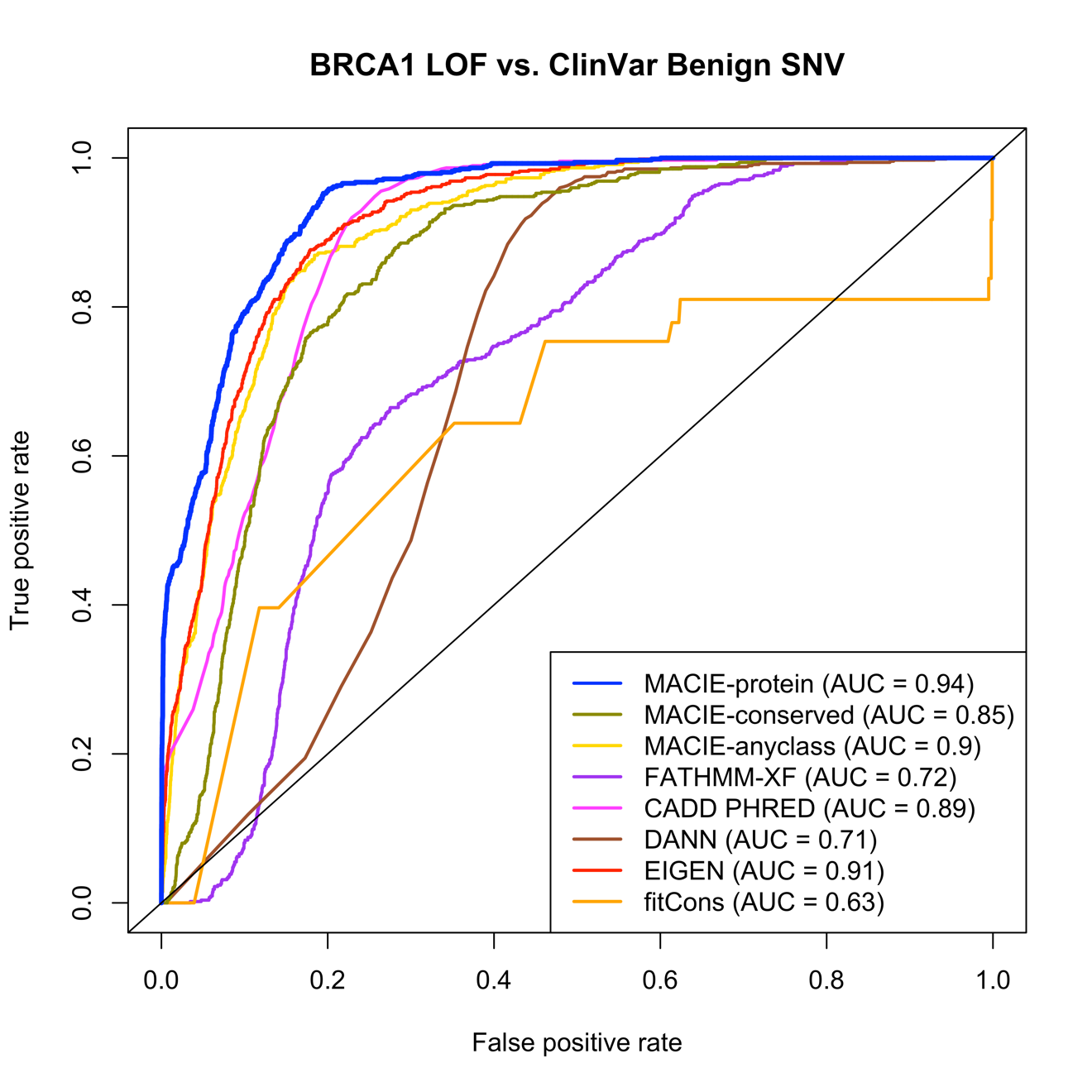

**Supplementary Figure 5.** LocusZoom plot for GWAS associations of TC at the *APOE* locus. The lipids GWAS summary statistics were from the European Network for Genetic and Genomic Epidemiology (ENGAGE) Consortium (*n* = 62,166). The MACIE-protein and MACIE-conserved scores for rs7412 are 0.96 and 0.97, respectively. The MACIE-conserved and MACIE-regulatory scores for rs1065853 are < 0.01 and > 0.99, respectively.

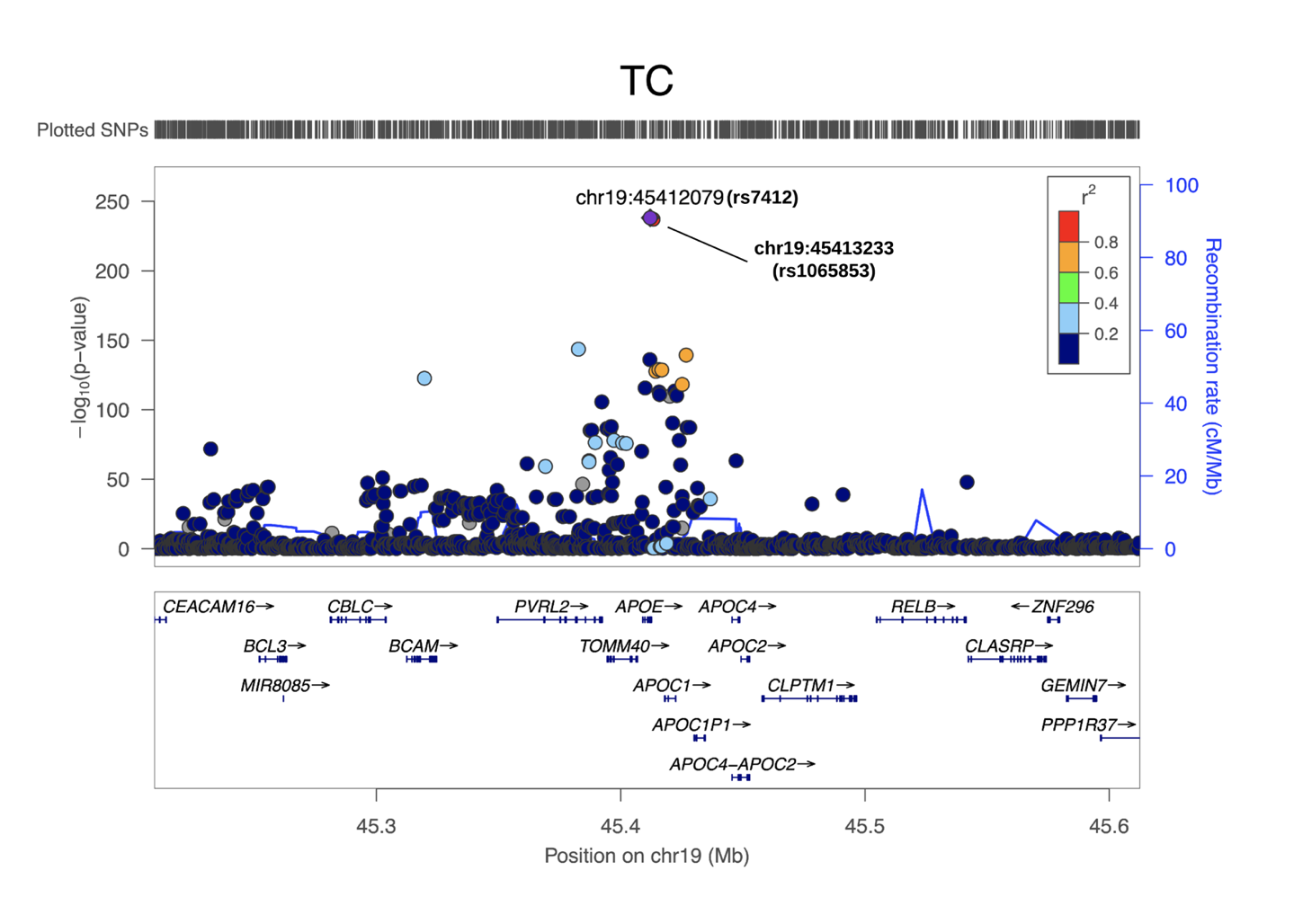

**Supplementary Figure 6.** LocusZoom plot for GWAS associations of HDL-C at the *CETP* locus. The lipids GWAS summary statistics were from the European Network for Genetic and Genomic Epidemiology (ENGAGE) Consortium (*n* = 60,812). The MACIE-conserved and MACIE-regulatory scores for rs17231506 are both < 0.01. For both rs72786786 and rs12720926, the MACIE-conserved and MACIE-regulatory scores are < 0.01 and > 0.99, respectively.

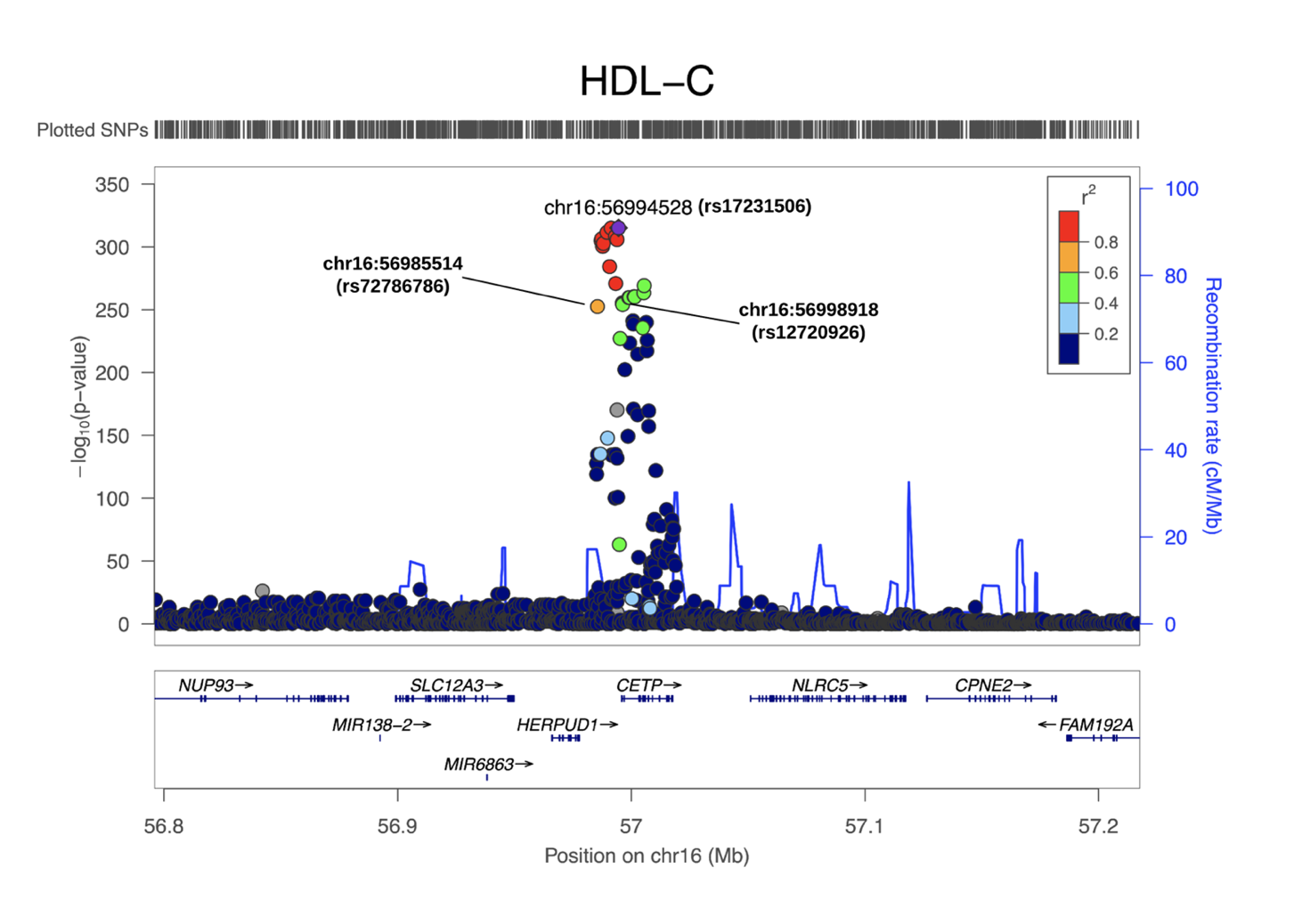

**Supplementary Figure 7.** LocusZoom plot for GWAS associations of TG at the *APOC3* locus. The lipids GWAS summary statistics were from the European Network for Genetic and Genomic Epidemiology (ENGAGE) Consortium (*n* = 60,027). The MACIE-conserved and MACIE-regulatory scores for rs964184 are both < 0.01. The MACIE-conserved and MACIE-regulatory scores for rs2075290 are < 0.01 and 0.88, respectively.

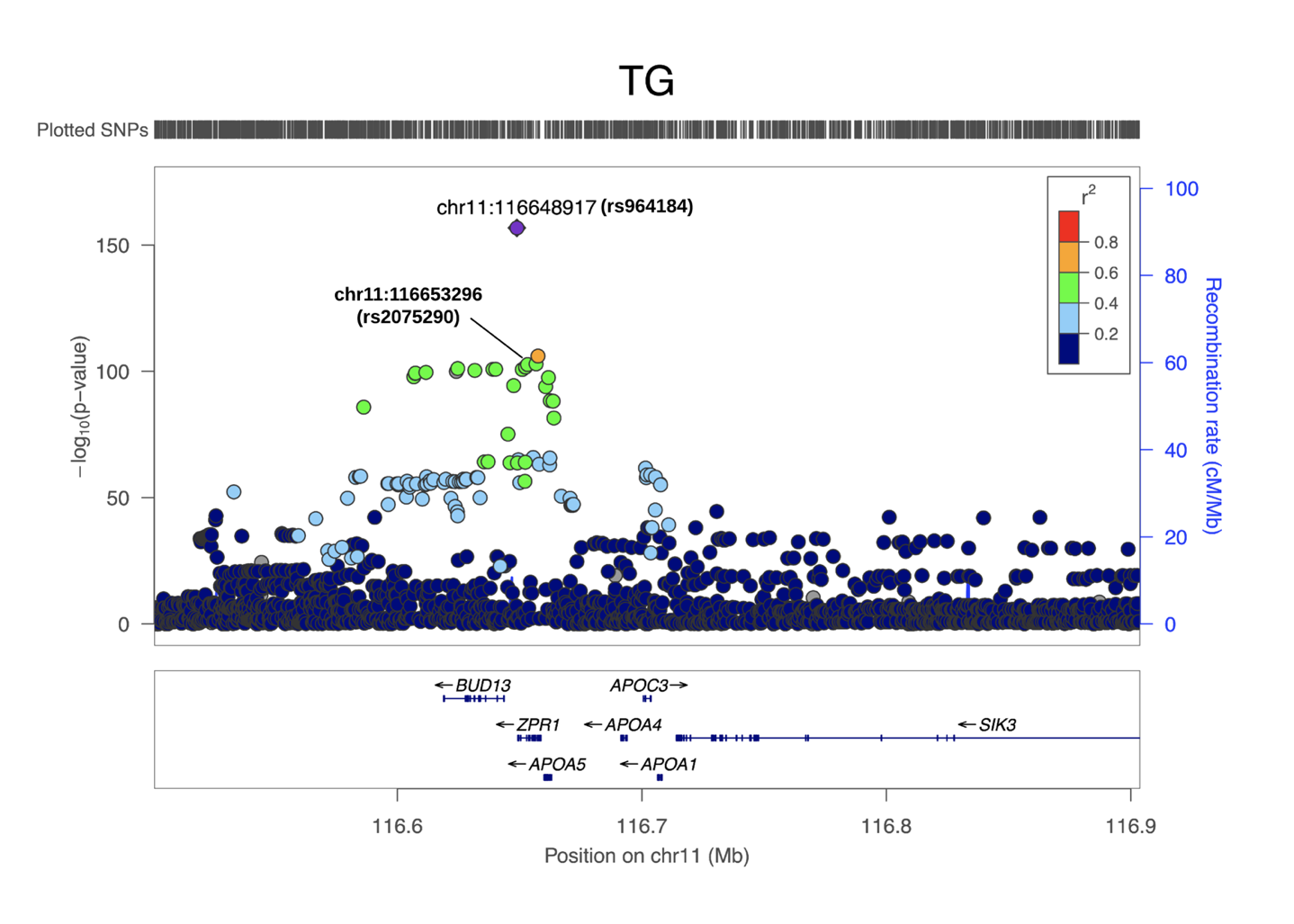
